# Seasonal biogeochemical variations in a modern microbialite reef under early Earth-like conditions

**DOI:** 10.1101/2025.02.22.639630

**Authors:** Federico A. Vignale, Laura Sánchez-García, Daniel Carrizo, Andrea Castillejos Sepúlveda, Heidi Taubner, Sebastián Oriolo, Alex L. Mitchell, Adrián G. Turjanski, Judith M. Klatt, Robert D. Finn, Maria M. Garcia-Alai, María E. Farías

**Affiliations:** European Molecular Biology Laboratory - Hamburg Unit, Notkestrasse 85, 22607 Hamburg, Germany; Centre for Structural Systems Biology, Hamburg, Germany; Centro de Astrobiología (CAB), CSIC-INTA, Torrejón de Ardoz, Madrid, 28850, Spain; Microsensor Group, Max Planck Institute for Marine Microbiology, Celsiusstraße 1, 28359 Bremen, Germany; MARUM, Center for Marine Environmental Sciences & Faculty of Geosciences, University of Bremen, 28359 Bremen, Germany; CONICET-Universidad de Buenos Aires, Instituto de Geociencias Básicas, Aplicadas y Ambientales de Buenos Aires (IGEBA), Intendente Güiraldes 2160, C1428EHA, Buenos Aires, Argentina; European Molecular Biology Laboratory, European Bioinformatics Institute (EMBL-EBI), Hinxton, CB10 1SD, UK; Laboratorio de Bioinformática Estructural, Instituto de Química Biológica de la Facultad de Ciencias Exactas y Naturales (IQUIBICEN)-CONICET, Universidad de Buenos Aires (UBA), Buenos Aires, Argentina; Microcosm Earth Center, University of Marburg & Max Planck Institute for Terrestrial Microbiology, 35032 Marburg, Germany; Biogeochemistry Group, Department of Chemistry, University of Marburg, Germany; PUNABIO S.A. Campus USP-T Av. Solano Vera y Camino a Villa Nougués, San Pablo, Tucumán, Argentina

**Keywords:** Andean lakes, microbial mats, microbialites, extremophiles, primitive Earth

## Abstract

Microbialites are organosedimentary structures dating to the Precambrian that serve as archives of Earth’s environmental evolution. Today, they persist in only a few environments markedly different from those in which they first arose. Here, we report a modern microbialite reef in Laguna Pozo Bravo (Puna region, Argentina), exposed to high radiation, low oxygen pressure, and volcanic inputs reminiscent of early Earth. Through physicochemical, mineralogical, spectroscopic, electron microscopy, and metagenomic analyses, we identified diverse microbial communities with metabolic capacities that induce mineralisation. Seasonal environmental fluctuations drive cyclical changes in community composition, producing potential mineralisation patterns. Our findings suggest that carbon fixation and the metabolic drivers of alkalinity in microbialites evolved over time. Moreover, the variability in prokaryotic compositions among modern microbialites demonstrates that carbonate precipitation is governed by metabolic potential rather than taxonomy, reinforcing their role as dynamic records of environmental conditions.

## Introduction

The Altiplano-Puna plateau, located in the central Andes mountains of South America, is comprised of numerous wetlands and salt flats. Due to the high altitude and volcanic activity of the area, these environments present extreme conditions. These include low oxygen pressure, intense solar radiation, low rates of precipitation, high rates of evaporation, pronounced seasonal and daily temperature fluctuations, strong winds, and elevated arsenic concentration^1^. Together, these factors make the region a valuable model for primitive Earth^2,3^. Some of these environments also share similarities with Mars, like the hypersaline lagoons resembling Martian paleolakes from the Noachian period (around 3,500 Ma)^4,5^, or the hydrothermal fields with widespread sinter deposits^6^, reminiscent of the opaline silica outcrops found in an ancient volcanic hydrothermal setting in Gusev crater^7^. Consequently, the study of the microbial ecosystems inhabiting these unique environments could provide valuable insights into the adaptations and survival strategies of early complex life on Earth, as well as refine our search for extraterrestrial life by identifying environmental signatures and biochemical markers indicative of life on other planets or moons^8^.

The microbial ecosystems of the Altiplano-Puna are complex associations of microorganisms with minerals, mainly halite (NaCl), gypsum (CaSO_4_-2H_2_O), calcite (CaCO_3_), and aragonite (CaCO_3_)^9^. These ecosystems can be classified, based on community organisation and extent of mineralisation, into biofilms, microbial mats, and microbialites, among others^1^. Biofilms are microbial communities attached to a surface forming a matrix of extracellular polymeric substances (EPS)^10^ that binds the cells together, protecting them from adverse conditions and facilitating their survival^11^. Microbial mats are stratified biofilms that can grow up to tens of centimetres thick. The microbial mat layers comprise species from different microbial guilds whose activity is governed by the availability of light, oxygen, and other resources^12^. The combination of microbial metabolism and nutrient transport, regulated by diffusion, generates steep concentration gradients of nutrients and metabolites which create distinct niches at various depth intervals^13^. Microbial mats have been around for more than 3,500 My^14^; however, the evolution of predatory metazoans and competition with macrophytes caused their decline 1,000 Ma. Today, these ecosystems develop only in aquatic environments where environmental adversities limit predation and competition. Such conditions are primarily found in hypersaline or geothermal habitats, like those in the Altiplano-Puna region^9,15^.

Some biofilms and microbial mats have the capacity to lithify, forming organosedimentary structures known as microbialites^16^. For this process to happen, two components must be present, first, an “alkalinity” engine and, second, an organic matrix comprised of extracellular polymeric substance (EPS) that provides a template for carbonate nucleation. The alkalinity engine is any process that can increase carbonate alkalinity, indirectly promoting carbonate precipitation. In other words, the alkalinity engine is any process that increases the calcium carbonate saturation index (SI), favouring precipitation. When the alkalinity engine is driven by microbial metabolic activities, the mineralization is biologically-induced. On the contrary, when the alkalinity engine is driven by environmental conditions, the mineralization is biologically-influenced. In the biologically-induced mineralization, photosynthesis and sulphate reduction are the microbial metabolic activities that increase the pH promoting carbonate precipitation, whereas aerobic respiration, fermentation, and sulphide oxidation are the metabolisms that decrease the pH triggering carbonate dissolution. The net carbonate precipitation will depend on the balance between these metabolic activities as well as their temporal and spatial variations. In the biologically-influenced mineralization, two main physicochemical processes lead to carbonate precipitation: evaporation of water and carbon dioxide degassing. In both types of mineralization, the organic EPS matrix serves as a template for mineral nucleation, but only when specific intrinsic conditions are met. The matrix contains abundant functional groups (hydroxyl and/or carboxyl groups) that bind Ca^2+^ or Mg^2+^ ions, inhibiting the precipitation of carbonate minerals. Therefore, to enable calcium carbonate precipitation, the Ca-binding capacity of the EPS matrix must be reduced. This can be achieved through EPS decomposition, alteration, or saturation with Ca^2+^ ^17,18^.

Microbialites can be classified as stromatolites^19^, thrombolites^20^, dendrolites^21^, and leiolites^22^ according to their laminated, clotted, shrub-like, and structureless mesostructures, respectively^23^. In general, microbialites are carbonatic in composition, although some are siliceous or even evaporitic^24^. The different types of microbialites result from different types of biofilms and microbial mats^25^, along with changes in environmental conditions^26,27^. Microbialites extend in the geological record from the Precambrian (∼3,500 Ma) to the present. Throughout Earth’s history, they experienced periods of expansion (between 2,800 and 1,000 Ma) and distinct periods of decline at approximately 2,000 Ma, 1,000 Ma, and 675 Ma, driven by major environmental and ecological changes^24,28–31^. Today, modern microbialites are distributed in few marine and continental environments, like those found in Shark Bay (Australia), Highborne Cay (The Bahamas)^32^, or the Altiplano-Puna (Argentina and Bolivia)^9^.

Over the past decade, explorations have been conducted in the Altiplano-Puna^9^ to identify and study extreme microbial ecosystems. Although numerous microbial mats and microbialites have been reported in the region^1^, the majority of these ecosystems remain undiscovered or poorly studied. In this work, we present an in-depth biogeochemical characterisation of a modern microbialite reef in Laguna Pozo Bravo, Salar de Antofalla, Argentina; which resembles conditions of ancient Earth. Our approach combined physicochemical analyses, mineralogy, bulk geochemistry, scanning micro X-ray fluorescence (µXRF), Raman spectroscopy, scanning electron microscopy (SEM), 16S rRNA (16S ribosomal RNA)) amplicon sequencing, and metagenomic analyses. This integrative analysis reveals the adaptation to seasonal fluctuations and metabolic functioning of the microbial communities inhabiting this unique ecosystem, as well as their role in mineral precipitation processes and biogeochemical cycles. We believe that these findings are vital for reconstructing the geobiological processes that shaped ancient sedimentary deposits on Earth.

## Results and discussions

### Laguna Pozo Bravo presents physicochemical conditions reminiscent of early Earth

Laguna Pozo Bravo (Fig. 1a) is a small (500 m long, 52 m wide) shallow (∼2.5 m deep) lagoon located at the foot of basaltic lava deposits in the northwest region of Salar de Antofalla (25° 30’ 49.82’’ S, 67° 34’ 38.92” W). Certain physicochemical conditions of this lagoon align with those of primitive Earth, as well as with Martian environments. Due to its high altitude, surface pressure at Pozo Bravo remains consistently low (613.9-618.2 hPa), approximately 60% of average sea level pressure^33^ (Supplementary Fig. S1). This reduced pressure leads to a partial pressure of oxygen between 128.9 and 129.8 hPa, considerably lower than the ∼210 hPa at sea level, and comparable to oxygen levels estimated for the late Paleoproterozoic to Mesoproterozoic (∼1,800–1,000 Ma) during the Precambrian^2,34^. This period marked a transitional phase in Earth’s oxygenation, with levels sufficient to sustain microbial mats and early eukaryotic life but still below those required for the emergence of complex multicellular organisms. Thus, Pozo Bravo serves as a model for Earth’s oxygen conditions before the evolution of complex multicellular life. The UV index is persistently elevated, with high values (∼6-9) in winter and extreme levels (∼17-19) in summer (Supplementary Fig. S2). These values are comparable to those expected on early Precambrian Earth during the Archean Eon (∼4,000–2,500 Ma), when a limited ozone layer allowed high UV radiation to penetrate the surface^35^. Land surface temperatures at Pozo Bravo show substantial fluctuations between day and night (ΔT up to 49.8 °C in summer) and across seasons (Supplementary Fig. S3). This diurnal temperature variation is akin to the extreme temperature swings seen on Mars, where day-night differences can reach up to 104 °C^36^. The lake water is alkaline (pH ∼8), highly saline (over 35000 mg/L), and rich in elements of volcanic origin, including sulphate (1,920-1,840 mg/L), boron (10.5-152 mg/L), lithium (21.3-101 mg/L), arsenic (0.07-0.86 mg/L), and manganese (0.05-0.60 mg/L) (Table 1). This water chemistry shares certain similarities with the mineral-rich conditions of Precambrian Earth’s oceans^37–40^ and the inferred composition of ancient Martian water bodies^41^, which likely held high concentrations of sulphate, chloride, and other volcanic derived elements. Overall, these extreme physicochemical conditions make Laguna Pozo Bravo a compelling model for ancient Earth, offering insights into environmental resilience and the potential for life in similar extraterrestrial environments.

**Fig. 1:**
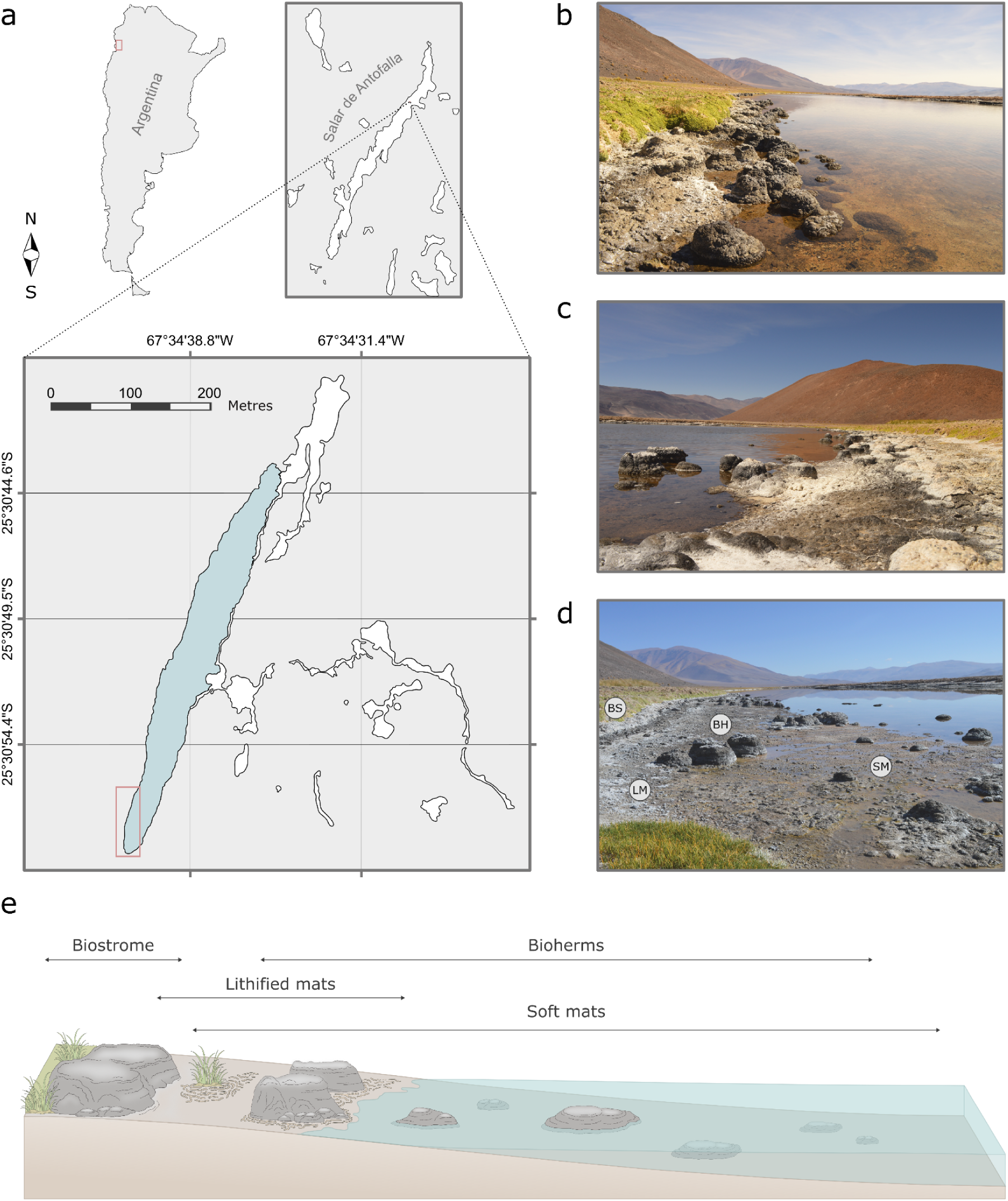
Laguna Pozo Bravo harbours a modern microbialite reef analogue for ancient Earth biostructures. **a** Location of Laguna Pozo Bravo in Salar de Antofalla, Catamarca (Argentina), and the sampling area. The sampling area (south-west margin of the lagoon) is highlighted with a red frame. **b** North view of the microbialite reef from the middle-west margin of the lagoon. **c** South view of the microbialite reef from the middle-west margin of the lagoon. **d** North view of the microbialite reef from the sampling area. **e** Transition of microbial structures along the intertidal zone in the sampling area. BS, Biostrome; LM, Lithified mats (carbonate pavement); BH, Bioherms; SM, Soft mats.

**Table 1.** Physicochemical parameters of surface water from Laguna Pozo Bravo.

| Physicochemical parameter | Unit | Winter value | Summer value |
| --- | --- | --- | --- |
| pH |  | 7.84 | 8 |
| Surface conductivity | mS/cm | 150.4 | 175.2 |
| Surface temperature | °C | 9 | 23 |
| Dissolved oxygen (DO) | % air saturation | 171 | 27.5 |
| Water column height of the shoreline | cm | 26 | 7 |
| Biochemical oxygen demand (BOD<sub>5</sub>) | mg/L | 161 | 230 |
| Chemical oxygen demand (COD) | mg/L | 475 | 705 |
| Total hardness | mg CaCO<sub>3</sub>/L | 8,460 | 10,250 |
| Chlorine | mg/L | 78,200 | 100,620 |
| Sodium | mg/L | 45,100 | 57,500 |
| Potassium | mg/L | 990 | 1,140 |
| Phosphate | mg/L | < 3.00 | < 3.00 |
| Nitrate | mg/L | < 5.00 | < 5.00 |
| Nitrite | mg/L | < 0.02 | < 0.02 |
| Sulphate | mg/L | 1,920 | 1,840 |
| Arsenic | mg/L | 0.07 | 0.86 |
| Calcium | mg/L | 1,570 | 1,910 |
| Lithium | mg/L | 21.30 | 101 |
| Magnesium | mg/L | 1,100 | 1,330 |
| Dissolved silica | mg/L | 83.20 | 158 |
| Iron | mg/L | < 0.10 | 0.33 |
| Boron | mg/L | 10.50 | 152 |
| Copper | mg/L | 0.24 | < 0.01 |
| Nickel | mg/L | < 0.05 | < 0.05 |
| Manganese | mg/L | 0.05 | 0.60 |
| Mercury | mg/L | < 0.01 | < 0.01 |
| Chromium | mg/L | < 0.01 | < 0.01 |
| Aluminium | mg/L | < 0.10 | 2.97 |
| Cadmium | mg/L | < 0.01 | < 0.01 |
| Lead | mg/L | < 0.01 | < 0.01 |

Laguna Pozo Bravo is a unique environment not only because it presents extreme physicochemical conditions, but also because these conditions vary between seasons (Table 1). In winter, when the water temperature is lower (9 °C) and the water level is higher (26 cm) due to reduced evaporation, the dissolved oxygen (DO) level is remarkably high at 171% air saturation. This high oxygen level corresponds with lower concentrations of ions (sodium, magnesium, chlorine, potassium, calcium) and minerals (e.g., Calcite and aragonite), resulting in lower conductivity (150.4 mS/cm) and total hardness (8,460 mg CaCO₃/L). In contrast, during summer, the elevated water temperature (23 °C) and lower water level (7 cm), driven by high evaporation rates, result in reduced dissolved oxygen (27.5% air saturation). The increased concentration of ions and minerals under these conditions leads to a rise in both conductivity (175.2 mS/cm) and total hardness (10,250 mg CaCO₃/L). Seasonal changes are also evident in biochemical oxygen demand (BOD5) and chemical oxygen demand (COD). Both BOD5 and COD values increase in summer to 230 mg/L and 705 mg/L, respectively, compared to their winter levels of 161 mg/L and 475 mg/L. This indicates a likely seasonal boost in microbial activity and biomass during the warmer months.

### Microbial mats in Pozo Bravo exhibit diel vertical geochemical gradients

The extreme conditions and minimal disturbance in Laguna Pozo Bravo create an environment that supports the growth of microbial mats, particularly along the lagoon’s southwest margin (Fig. 1a, d, e), where they exhibit cerebroid- or snake-like morphologies due to biogenic gas accumulation (Fig. 2a). In the absence of significant physical disruption, microbial mats organise into layers, each containing microorganisms with distinct metabolic activities. The microbial mats in Pozo Bravo display three main layers: a green surface layer, a reddish-pink middle layer, and a dark brown underlying layer (Fig. 2b).

**Fig. 2:**
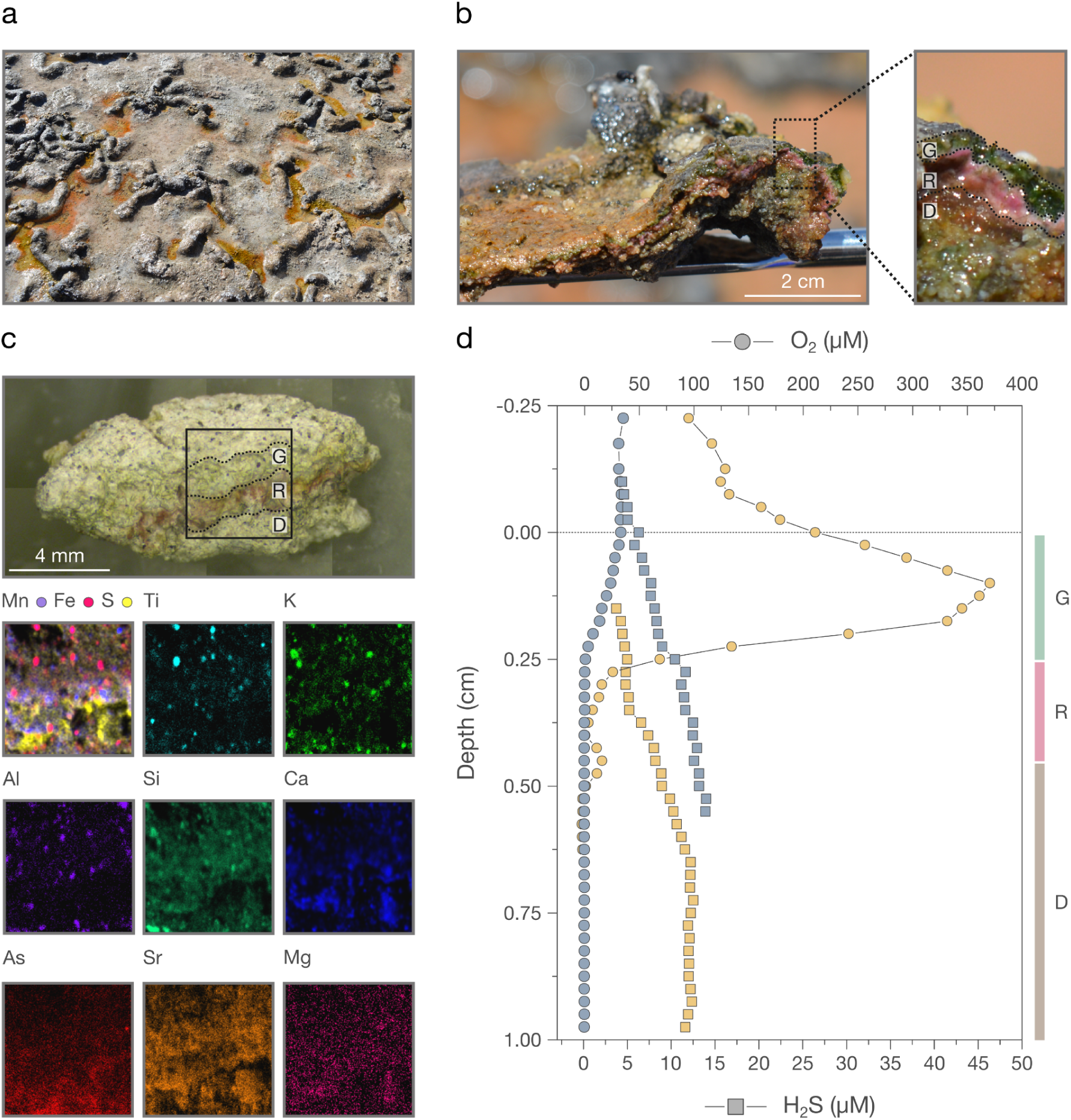
Pozo Bravo microbial mats show spatial and temporal heterogeneity in their chemical composition. **a** Overview of Pozo Bravo soft microbial mats displaying cerebroid- or snake-like structures. **b** Close-up look at a cross-section of a soft microbial mat exhibiting a green surface layer (G), a reddish-pink middle layer (R), and a dark brown underlying layer (D). **c** Spatial distribution of elements (deconvoluted counts) in a freeze-dried soft microbial mat sample determined by µXRF spectrometry. The area of µXRF scanning (4.2 mm x 4.3 mm) is marked with a black frame, and the dotted lines highlight the green, reddish-pink, and dark brown layers. **d** Oxygen and sulphide profiles (mean values, N=3) of a soft microbial mat, measured *in situ* during the middle of the day (from 13:00 to 16:00; yellow symbols) and night (from 20:00 to 01:00; blue symbols).

Microbial mat layered structures are shaped by steep vertical geochemical gradients generated by environmental factors and the metabolic activities of mat organisms. In the Pozo Bravo microbial mats, oxygen is detectable in the green surface layer, where its concentration rises during the day (370.69 µM at 0.1 cm depth) and drops at night (24.00 µM at 0.1 cm depth) (Fig. 2d), suggesting oxygenic phototrophic activity. With oxygen available mainly in this layer, aerobic respiration is primarily confined to it. Manganese is abundant between the green and reddish-pink layers (Fig. 2c), suggesting active manganese oxidoreduction in this zone. Sulphur is concentrated within the reddish-pink and dark brown layers (Fig. 2c), which correlates with the observed sulphide profiles (Fig. 2d). In the reddish-pink layer, sulphide levels decrease during the day (4.80 µM at 0.3 cm depth) but increase again at night (11.88 µM at 0.3 cm depth), indicating anoxygenic phototrophic activity driven by sulphide oxidation. The fact that sulphur concentrates in the dark brown layer further suggests sulphur oxidoreduction activity in this area (Fig. 2c). Arsenic also accumulates in the dark brown layer, implying potential arsenic oxidoreduction activity within this anaerobic region (Fig. 2c).

The observed spatial and temporal variations in chemical composition suggest distinct microbial communities within each mat layer (Fig. 2). The green surface layer is dominated by oxygenic phototrophs, which couple inorganic carbon assimilation to light energy, oxidising water and producing oxygen as a byproduct. This layer thereby provides an ideal micro-environment for aerobic and facultative aerobic heterotrophs. Aerobic heterotrophs gain energy by oxidising organic carbon exudates through oxygen respiration, while facultative aerobic heterotrophs can switch to using other terminal electron acceptors when oxygen is limited. The reddish-pink layer is dominated by anoxygenic phototrophs, which use sulphide as an electron donor for photosynthesis and release sulphate as a byproduct. The dark brown layer is dominated by anaerobic heterotrophs and fermenters. Anaerobic heterotrophs, mainly sulphate reducing bacteria (SRB), oxidise organic carbon using a range of terminal electron acceptors, while fermenters use organic carbon as both electron donor and acceptor. Sulphide oxidising bacteria, which oxidise reduced sulphur compounds with oxygen or nitrate, might also be abundant in the dark brown layer.

### Both eukaryotes and prokaryotes participate in the organomineralization process

In Laguna Pozo Bravo, the formation of lithified mats and microbialites (Fig. 1b-e) is driven by the constantly elevated calcium carbonate saturation index (aragonite SI is 2.78 in winter and 3.36 in summer; calcite SI is 2.94 in winter and 3.54 in summer), which indicates supersaturation of this mineral throughout the year, as well as the presence of abundant EPS, which serve as a template for carbonate nucleation (Fig. 3). The elevated calcium carbonate saturation index (Supplementary Tables S2 and S3) likely results from environmental conditions like water evaporation, reflected in the seasonal variations in water column height (Table 1). Nevertheless, the spatial and temporal separation of metabolic activities observed in the microbial mats (Fig. 2) may also be responsible for local differences in the saturation index which promote carbonate precipitation. Consequently, mineralization in Pozo Bravo may be both biologically induced and influenced. Chemical analyses conducted on the microbial mats revealed increased concentrations of arsenic, calcium, magnesium, iron, boron, and manganese compared to their levels in the water (Table 2 and Supplementary Table S1). This suggests that the EPS matrix of the mats acts as a chelator for cations. Therefore, the precipitation of calcium carbonate is likely enhanced by mechanisms that weaken the capacity of the EPS matrix to bind cations. For instance, extreme UV radiation (Supplementary Fig. S2) may contribute to partial EPS destruction through Maillard browning reactions (involving the reaction of sugars and aminoacids), whereas the activity of heterotrophic microorganisms may lead to microbial decomposition of the EPS. Additionally, the notably higher concentrations of calcium and magnesium in the mats compared to the water suggest that saturation of the EPS cation-binding capacity might also facilitate calcium carbonate precipitation^18^.

**Fig. 3:**
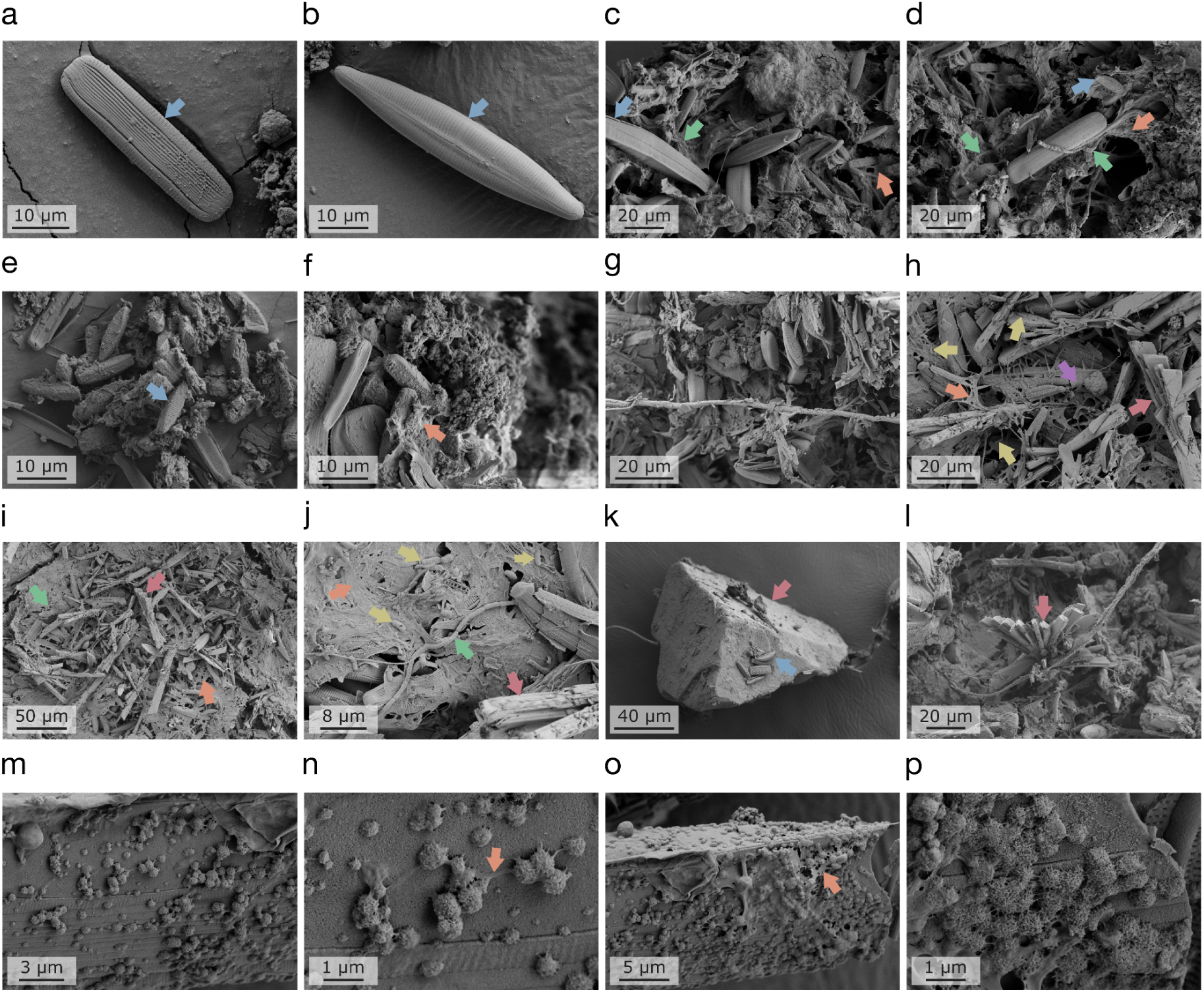
Diatoms, cyanobacteria, and prokaryotic cells participate in the lithification process of Pozo Bravo microbial mats. **a-p** Scanning electron microscopy images (SEM) from microbial mats samples. Different recognized structures are marked with an arrow: Orange, EPS serving as a template for carbonate precipitation; Red, mineral structure; Green, filamentous cyanobacterium; Yellow, bacillus; Purple, spirochete; Blue, diatom.

**Table 2.** Physicochemical parameters of a soft microbial mat from Laguna Pozo Bravo.

| Physicochemical parameter | Unit | Winter value | Summer value |
| --- | --- | --- | --- |
| pH |  | 7.3 | 7.6 |
| Conductivity 1:5 | mS/cm | > 20 | 8.32 |
| Humidity | % w/w | 58.6 | 47.3 |
| Organic material | % w/w | 10.4 | 13.3 |
| Chlorine | mg/Kg | 101,000 | 25,620 |
| Sodium | mg/Kg | 66,900 | 24,800 |
| Potassium | mg/Kg | 22,500 | 1,040 |
| Phosphate | mg/Kg | < 50.00 | < 50.00 |
| Total phosphorus | mg/Kg | 86.7 | 173 |
| Nitrate | mg/Kg | < 1.00 | < 50.00 |
| Nitrite | mg/Kg | 390 | < 1.00 |
| Ammoniacal nitrogen | mg/Kg | 140 | 457 |
| Sulphate | mg/Kg | 2,720 | 1,690 |
| Arsenic | mg/Kg | 282 | 201 |
| Calcium | mg/Kg | 121,000 | 144,000 |
| Lithium | mg/Kg | 145 | 58.6 |
| Magnesium | mg/Kg | 13,000 | 14,516 |
| Dissolved silica | mg/Kg | 2,200 | 994 |
| Iron | mg/Kg | 5,780 | 1,621 |
| Boron | mg/Kg | 1,100 | 371 |
| Copper | mg/Kg | 32.5 | 13.3 |
| Nickel | mg/Kg | < 5.00 | 1.7 |
| Manganese | mg/Kg | 3,760 | 1,996 |
| Bromine | mg/Kg | < 50.00 | < 50.00 |
| Mercury | mg/Kg | < 0.80 | < 0.10 |
| Chromium | mg/Kg | 52.6 | 1.3 |
| Aluminium | mg/Kg | 6,580 | 1,547 |
| Cadmium | mg/Kg | 17.3 | 1.1 |
| Lead | mg/Kg | < 20.0 | < 0.60 |
| Tin | mg/Kg | < 50.00 | < 1.25 |
| Cyanide | mg/Kg | < 50.00 | < 0.25 |

The described mineralization process has been further analysed through scanning electron microscopy (SEM) of a microbial mat sample (Fig. 3). The SEM images obtained revealed the presence of diatoms (*Bacillariophyceae*), filamentous cyanobacteria, and other prokaryotic cells (cocci, bacilli, and spirilla) forming aggregates embedded on the EPS matrix, where abundant carbonate precipitation takes place. This suggests the involvement of both prokaryotic and eukaryotic microorganisms in the mineral precipitation processes that leads to the formation of a lithified microbial mat or microbialite. Based on morphological characteristics, we could identify at least three genera of diatoms: *Navicula* (Fig. 3a and b), *Halamphora* (Fig. 3c, d, and k), and *Nitzschia* (Fig. 3e), previously reported to compose microbial mats and microbialites in the central Andes region^9^.

### Environmental conditions and microbial activity shape microbialite structure and composition

In Laguna Pozo Bravo, modern microbialites extend along the entire margin, forming a reef analogue for early Earth biostructures (Fig. 1b-e). These microbialites vary in height from 0.05 to 0.65 m and display a wide range of macrostructures, including domical, discoidal, and tabular shapes (Fig. 4a). However, the most distinctive feature of these microbialites is their internal organization, which transitions gradually from a thrombolite core to a dendrolite middle layer and, finally, to a stromatolite surface^42^ (Fig. 4b). Thrombolite and dendrolite mesostructures are characterized by lighter colouration, whereas stromatolite mesostructures appear darker (Fig. 4b and c). Consequently, we have divided the internal organization of the microbialites into two distinct zones: (I) an internal light area and (II) an external black rim. Notably, these two zones are also present in the lithified microbial mats, suggesting that microbial mats and microbialites undergo similar mineralization processes.

**Fig. 4:**
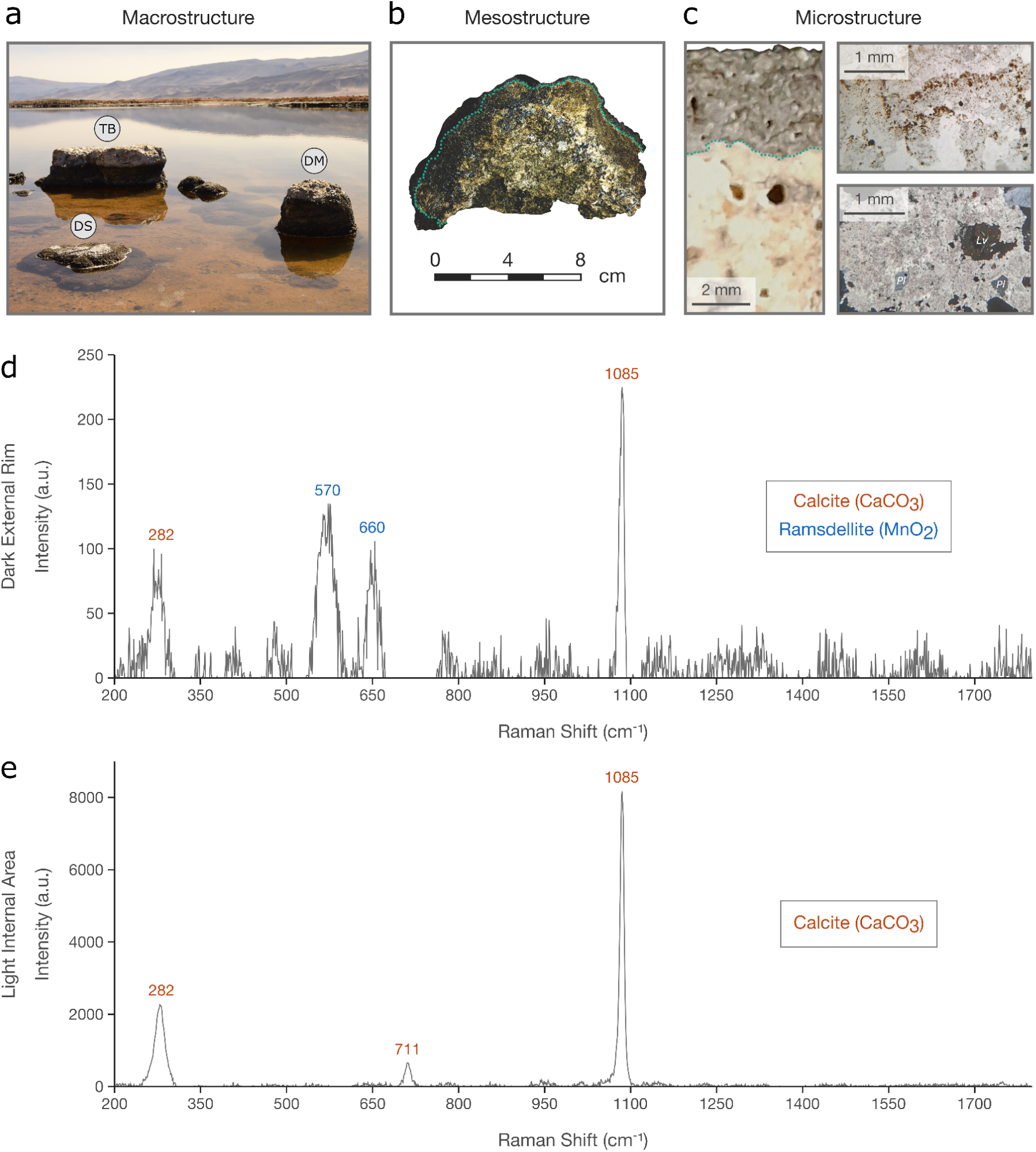
Pozo Bravo microbialites are morphologically, microstructurally and mineralogically heterogeneous. **a** Overview of Pozo Bravo bioherms displaying diverse external morphologies (macrostructure). DM, domical shape; DS, discoidal shape; TB, tabular shape. **b** Cross-section of domical bioherm, showing an external dark rim followed by an internal light area. The division between the two areas is indicated by a green dotted line. **c** On the left, a polished hand specimen of a domical bioherm showing the dark external rim (above) and the light internal area (below). On the top-right, a photomicrograph of the dark external rim (plain-polarised light) with thin layers of dark reddish oxides, corresponding to manganese oxides intergrown with calcite. On the bottom-right, a photomicrograph of the light internal area (cross polars) dominated by calcite with detrital grains of plagioclase (Pl) and volcanic rock lithic fragments (Lv). **d** Representative Raman spectra in the dark external rim characterised by ramsdellite and calcite peaks. **e** Representative Raman spectra in the light internal area characterised by calcite bands.

A Raman spectroscopy analysis of a microbialite sample (Fig. 4d and e) revealed that both distinct internal zones are primarily composed of calcite. However, the dark external rim also contains ramsdelite, a black manganese dioxide (MnO₂) mineral, which accounts for the observed colour differences between the two zones. The presence of ramsdellite in the outermost zone, which contributes to the external colouration of microbialites and lithified mats, can be attributed to microbial activity. Microorganisms, particularly manganese-oxidising bacteria, are well-known for their ability to mediate the precipitation and transformation of manganese minerals, including manganese oxides like ramsdellite^43,44^. The microbial oxidation of soluble Mn(II) ions in Pozo Bravo is supported by the high manganese levels in the water (Table 1) and the substantial manganese concentration found between the green and reddish-pink layers of the microbial mat sample (Fig. 2c), where the transition from oxic to anoxic conditions occurs. Geochemical evidence linking manganese cycling to microbial processes, such as that observed in Pozo Bravo, could prove useful in the search for potential life beyond our planet, particularly on Mars, where the Curiosity rover has detected high concentrations of manganese oxides in rocks from Gale Crater^45,46^.

Powder X-Ray diffraction analyses of microbial mat (soft and lithified) and microbialite samples (Supplementary Table S4) further demonstrated how environmental conditions and microbial activities shape the mineralogical composition of these ecosystems. Both types of microbial mats are predominantly composed of magnesium calcite, with minor contributions from silicate minerals such as quartz (SiO_2_) and plagioclase feldspars. In the soft mats, the plagioclase component is primarily andesine ((Ca,Na)(Al,Si)_4_O_8_), while the lithified mats contain albite (NaAlSi_3_O_8_) and anorthite (CaAl_2_Si_2_O_8_). In contrast, microbialites are primarily composed of calcite throughout their structure, with minor contributions from hydrotalcite (Mg_6_Al_2_CO_3_(OH)_16_·4H_2_O), illite ((K,H_3_O)(Al,Mg,Fe)_2_(Si,Al)_4_O_10_[(OH)_2_,(H_2_O)]), and silicate minerals, including quartz, albite, andesine, and labradorite ((Ca,Na)(Al,Si)_4_O_8_). Microscopic examination of the microbialite samples (Fig. 4c) suggests that quartz and plagioclase are detrital clasts, likely sourced from adjacent volcanic rocks. This volcanic input may also account for the presence of hydrotalcite, which is commonly formed as an alteration product of basalts in alkaline water environments^47^. The presence of low-magnesium calcite in the microbialites may also be the result of post-depositional alteration, where magnesium ions in magnesium calcite are replaced by calcium ions, leading to the formation of a more stable mineral. These findings indicate that the mineralogical composition of microbial mats and microbialites reflects a combination of biogenic and detrital inputs, with microbialites further shaped by diagenetic transformations.

Bulk geochemical analyses of soft microbial mat and microbialite samples (Supplementary Table 5) revealed notable differences in the content of total organic carbon (TOC) and total nitrogen (TN). The soft microbial mat sample exhibited higher levels of TOC (2.1% of dry weight) and TN (0.42%), compared to the microbialite sample, which had only 0.35% TOC and no detectable TN. This suggests that microbial mats exhibit greater microbial activity and organic matter accumulation than microbialites. The bulk carbon isotopic composition (δ^13^C) of both the soft microbial mat (-18.2‰) and microbialite (-23.4‰) samples indicate carbon fixation through the reductive pentose phosphate cycle (Calvin−Benson−Bassham cycle), which is employed by oxygenic (e.g. diatoms and cyanobacteria) and anoxygenic (e.g. purple sulphur bacteria) phototrophs^48–51^. The bulk nitrogen isotopic composition (δ^15^N) of the soft microbial mat sample (0.4 ‰) indicates nitrogen fixation, likely by cyanobacteria, through nitrogenase enzymes^52^. These results are consistent with the phototrophic activities inferred from the microelectrode measurements (Fig. 2d) and the organisms identified in the SEM images (Fig. 3) within the microbial mats.

### The microbialite reef in Pozo Bravo shows seasonal variations in microbial community composition

A molecular diversity analysis based on the small subunit rRNA gene of *Bacteria* and *Archaea* was conducted over an annual cycle to study the prokaryotic composition of the water, microbial mats, and microbialites (Fig. 5). The results revealed significant differences in community composition (PERMANOVA, R² = 0.608, *p* ≤ 0.001) between the water and the benthic microbial ecosystems (Figs. 5a and c). The microbial community in the water is dominated by two phyla, *Bacteroidota* (60.91–88.31%) and *Pseudomonadota* (11.10–37.69%), largely represented by the classes *Flavobacteriia* (60.49–88.12%) and *Gammaproteobacteria* (10.40–35.83%). These classes are composed mainly of heterotrophic and halophilic bacteria from the genera *Psychroflexus*, *Owenweeksia*, and *Halomonas*. The prevalence of these taxa reflects notable organic matter processing in the water and indicates strong adaptations to the hypersaline conditions and elevated organic content of the environment. In contrast, microbial mats and microbialites support more diverse and complex microbial communities (Fig. 5b). These communities are dominated by several phyla, including *Pseudomonadota* (24.34-63.67%), *Bacteroidota* (3.42-38.35%), *Bacillota* (0.18-23.64%), *Cyanobacteriota* (1.02-14.31%), *Spirochaetota* (0.02-12.21%), *Chloroflexota* (0.53-8.29%), *Deinococcota* (0.25-5.49%), *Planctomycetota* (1.44-4.46%), and *Verrucomicrobiota* (0.08-2.93%). *Pseudomonadota* is mainly represented by the classes *Gammaproteobacteria* (4.52-56.02%), *Alphaproteobacteria* (6.64-20.70%), and *Deltaproteobacteria* (0.64-10.87%). These groups include anoxygenic phototrophs such as purple sulphur bacteria (*Thiocapsa*, *Thiococcus, Thiocystis*, *Thioflavicoccus*, *Thiohalocapsa*, *Thiorhodococcus, Thiorhodovibrio*, *Halochromatium*, *Ectothiorhodospira*, and *Halorhodospira*) and purple non-sulphur bacteria (*Rhodomicrobium*, *Roseospira*, *Rhodospira*, and *Rhodovibrio*), heterotrophic sulphur oxidisers (*Sulfurimonas*, *Sulfurifustis*, *Sedimenticola*, *Desulfobacula*, *Desulfobacter*, *Desulfovibrio*, and *Desulfuromonas*), sulphur reducers (*Desulfobacter*, *Desulfobacula*, *Desulfovibrio*, *Desulfuromonas*, *Desulfuromusa*, *Shewanella*, and *Geobacter*), and metal/metalloid reducers (*Desulfuromonas*, *Desulfuromusa*, *Desulfovibrio*, *Geobacter*, and *Shewanella*) capable of reducing iron, arsenic, manganese, cobalt, selenium, or chromium. The phyla *Bacteroidota, Bacillota*, *Spirochaetota*, *Deinococcota*, *Planctomycetota*, and *Verrucomicrobiota* are primarily represented by the classes *Bacteroidia* (0.11-10.61%), *Cytophagia* (0.48-8.41%), *Flavobacteriia* (0.62-18.84%), *Saprospiria* (0.20-8.17%), *Deinococci* (0.25-5.48), *Bacilli* (0.00-23.08%), *Clostridia* (0.12-12.19%), *Phycisphaerae* (0.30-2.68%), *Spirochaetia* (0.02-12.19%), and *Opitutae* (0.00-1.24%). These groups are predominantly composed of heterotrophic bacteria, with some capable of sulphur reduction (*Bacteroides*, *Marinifilum*, *Anaerophaga*, *Clostridium, Desulfotomaculum*, *Halanaerobium*, *Spirochaeta*, and *Sediminispirochaeta*), sulphur oxidation (*Cytophaga*, *Clostridium*, *Flavobacterium*, and *Spirochaeta*), ammonification (*Bacteroides* and *Anaerophaga*), nitrite reduction (*Flavobacterium* and *Maribacter*), and denitrification or nitrogen fixation (*Clostridium*, *Desulfotomaculum*, *Halanaerobium*, and *Bacillus*). *Cyanobacteriota* is represented by oxygenic phototrophs from different genera, such as *Halothece*, *Coleofasciculus*, *Halomicronema*, and *Synechococcus*. Among these genera, *Coleofasciculus* and *Synechococcus* include some species capable of nitrogen fixation. Meanwhile, *Chloroflexota*, primarily represented by *Anaerolineae* (0.33–5.29%), includes green non-sulphur bacteria capable of anoxygenic photosynthesis (*Candidatus* Chlorothrix). The prokaryotic composition observed in the microbial mats and microbialites aligns with the microbial activities inferred from the microelectrode measurements (Fig. 2d) and the bulk carbon and nitrogen isotopic compositions (Supplementary Table S5). The diversity of bacterial genera capable of oxygenic and anoxygenic photosynthesis, nitrogen fixation, and sulphur and nitrogen oxidoreduction highlights the complex metabolic interactions driving the biogeochemical cycles within these benthic ecosystems. This diversity contrasts with that of the water column, which is dominated by a few heterotrophic taxa, reflecting the distinct ecological roles of these habitats.

**Fig. 5:**
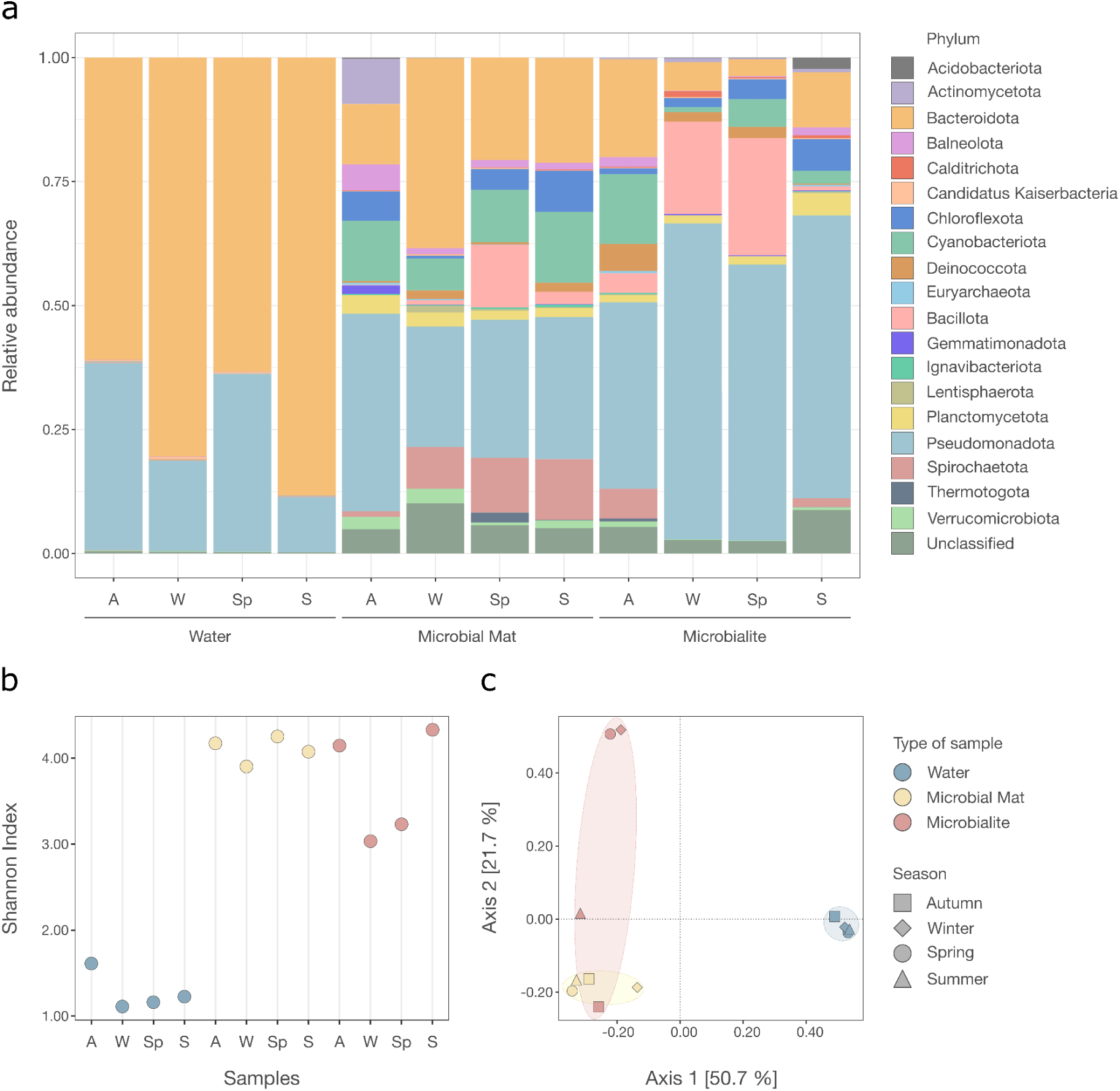
Prokaryotic diversity in the water, soft microbial mats, and microbialites changes throughout the seasons. **a** Relative abundance of the twenty major prokaryotic phyla in the different seasons. A, Autumn; W, Winter; Sp, Spring; S, Summer. **b** Comparison of alpha diversity metrics (Shannon index) between samples throughout the different seasons. **c** Principal coordinate analysis (PCoA) based on Bray-Curtis dissimilarity between samples throughout the different seasons.

The molecular diversity analysis also revealed seasonal variations in the prokaryotic composition of microbial mats and microbialites. Cyanobacteria increase in abundance during periods of higher light availability and decrease when light levels are lower (Fig. 5 and Supplementary Fig. S5). In the microbial mats, they reach their highest relative abundances in summer, whereas in the microbialites, the peaks occur in autumn and spring. These differences may be related to the greater susceptibility of microbialites to desiccation, making summer conditions more extreme and thus less favourable for cyanobacterial proliferation. Green non-sulphur bacteria show a similar pattern, peaking in summer and spring, reflecting their strong dependence on light availability as well. Purple sulphur bacteria also increase during months with higher light, but reach their maximum abundances primarily in autumn in both microbial mats and microbialites (Supplementary Fig. S5). This deviation from a mid-summer peak, as seen in cyanobacteria and green non-sulphur bacteria, may be linked to the activity of sulphur-reducing bacteria (SRB). SRB show greater abundances in spring and summer in the mats, and in autumn in the microbialites (Supplementary Fig. S5). Elevated levels of sulphide produced by SRB could promote the autumnal bloom of purple sulphur bacteria. In contrast, cyanobacteria and green non-sulphur bacteria do not depend on sulphide, which could explain why their seasonal trends align more directly with light intensity rather than the sulphur cycle. These findings indicate that multiple environmental factors, including light availability and the presence of redox-active species, drive the seasonal compositional shifts and metabolic interactions within these microbial ecosystems.

### Symbiotic microorganisms regulate biogeochemical cycles that shape the structure and mineralization of mats

A metagenomic analysis was performed on a soft microbial mat to explore the metabolic diversity of their microorganisms and to elucidate their roles in the biogeochemical cycles (Supplementary Table S6). The high relative abundance of genes encoding subunits of Photosystems I and II (*psa* and *psb*, respectively), along with the presence of *puf* genes associated with bacterial phototrophic reaction centers, suggests co-occurring oxygenic and anoxygenic photosynthesis within the microbial mat (Fig. 6a), as confirmed by microelectrode measurements. The presence of *nif* genes indicates that the microorganisms in the mat fix atmospheric nitrogen through nitrogenase enzymes, as reflected by the bulk nitrogen isotopic composition (δ^15^N=0.4‰) (Supplementary Table S5). Since *vnf* genes encoding vanadium-dependent nitrogenases were not identified, nitrogen fixation is likely carried out solely by molybdenum-dependent nitrogenases (*nif)*. Regarding carbon fixation, the high relative abundance of *cbb* and *prk* genes suggests that carbon fixation in the mat is primarily carried out through the reductive pentose phosphate cycle (Calvin–Benson–Bassham cycle), which is consistent with the bulk carbon isotopic composition results (δ¹³C=-18.2‰) (Supplementary Table S5). However, the presence of *acl* genes associated with the reductive tricarboxylic acid cycle 1 (rTCA cycle 1), *acs* genes involved in the reductive acetyl-CoA pathway (Wood–Ljungdahl pathway), and *mcl* and *mcr* bacterial genes related to the 3-hydroxypropionate bicycle also indicates carbon fixation through alternative pathways/cycles. Genes involved in the oxidation of hydrogen (*hya*), sulphur (*sqr*, *fcc*, *sox*, *dsr*, *hdr*, *sor*, *soe*, *apr*, *sat*, and *tcdh*), nitrogen (*hao* and *nxr*), arsenic (*aio*), and manganese (*mnxG*) compounds were also identified, indicating that the microorganisms in the mat are actively engaged in these processes. Some of them may even be obtaining energy through the oxidation of these compounds. Finally, the identification of genes involved in the reduction of sulphur (*sat*, *apr*, and *dsr*), nitrogen (*narG*, *napA*, *nir*, *norB*, and *nosZ*), iron (*mtr*), and arsenic (*arr*) compounds suggests that some microorganisms in the mat could use alternative electron acceptors to oxygen for respiration. In fact, the mtr system has been implicated not only in the reduction of iron but also in the reduction of manganese, cobalt, and chromium^53^, expanding the possibilities for anaerobic respiration in Pozo Bravo. These findings align with the microelectrode measurements, which revealed a large anoxic section in the microbial mat where anaerobic microorganisms flourish.

**Fig. 6:**
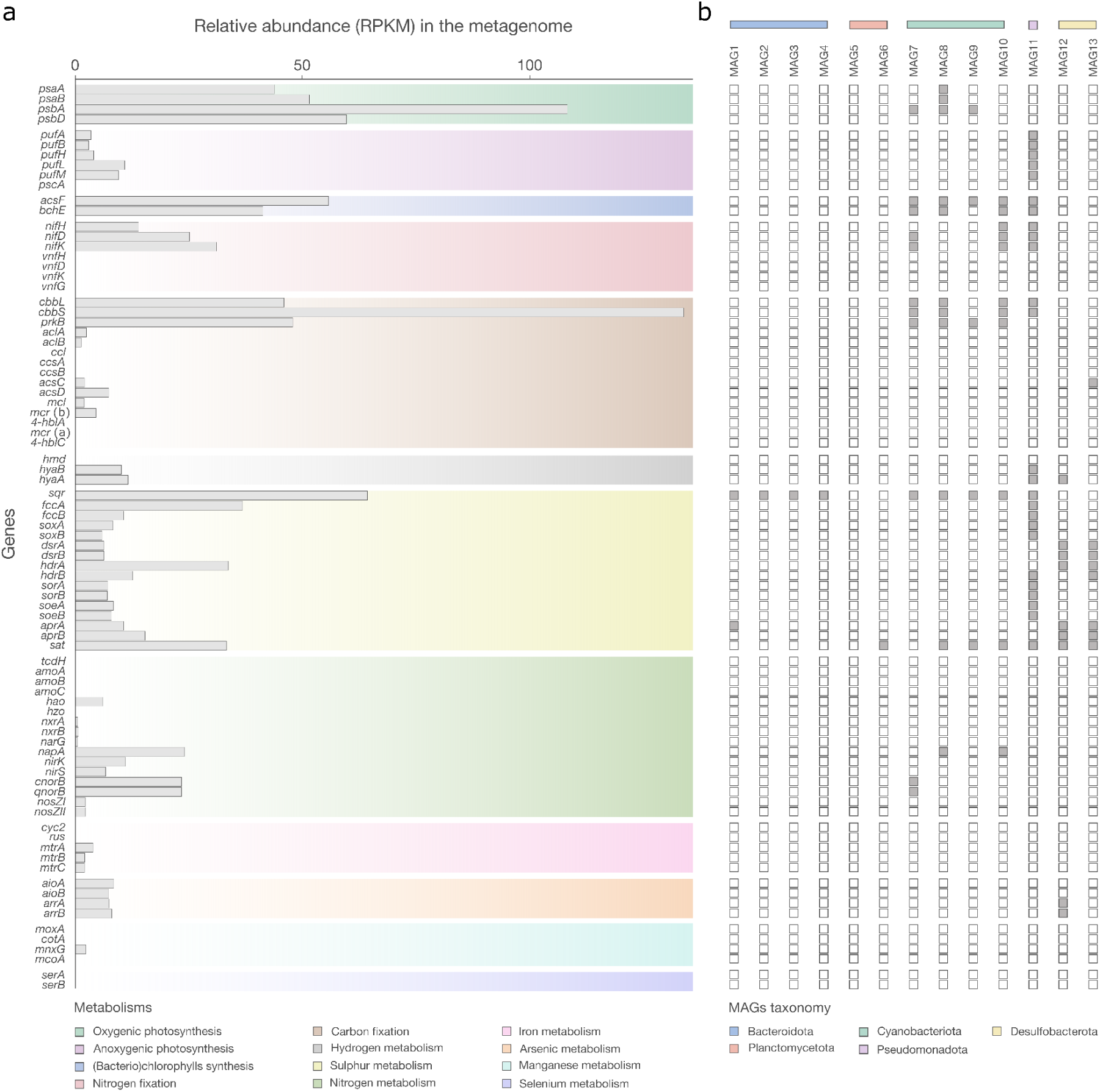
Microorganisms drive main biogeochemical cycles in Pozo Bravo’s soft microbial mats. **a** Relative abundance in the metagenome (presented as reads per kilobase per million mapped reads, RPKM) of genes associated with hydrogen, carbon, nitrogen, oxygen, sulphur, manganese, iron, arsenic, and selenium cycling. **b** Metabolic potential of the metagenome-assembled genomes (MAGs). Grey squares indicate the presence of a gene.

The reconstruction of thirteen metagenome-assembled genomes (MAGs) from the metagenomic contigs further expanded our understanding of the biogeochemical cycles in the microbial mats (Fig. 6b and Supplementary Table S7). Of these thirteen, four are nearly complete and nine are of medium quality. However, they are estimated to have very low contamination, with seven MAGs having contamination levels less than or equal to 1% (see Methods). Four of these MAGs belong to the phylum *Cyanobacteriota*, with two (MAGs 7 and 8) identified as members of the family *Geitlerinemaceae* and the other two (MAGs 9 and 10) classified under the genus *Halothece* (Supplementary Tables S8). MAGs 7 and 10 contain *nif*, *napA*, and *norB* genes, suggesting that cyanobacteria would play a key role in the nitrogen cycle, particularly in nitrogen fixation and denitrification. All four cyanobacterial MAGs possess genes for carbon fixation (*cbb* and *prk*) and hydrogen sulphide oxidation (*sqr*). On the one hand, the presence of *cbb* and *prk* genes indicates that cyanobacteria drive carbon fixation through the reductive pentose phosphate cycle, which is the primary carbon fixation process in the mat. On the other hand, the detection of the *sqr* gene suggests that cyanobacteria may use hydrogen sulphide oxidation as an alternative electron source for photosynthesis, resembling some anoxygenic phototrophs. The ability of cyanobacteria to perform anoxygenic photosynthesis via hydrogen sulphide oxidation is supported by the microelectrode measurements (Fig. 2d), which showed a decrease in the hydrogen sulphide levels during the day in the green surface layer, where cyanobacteria are found (Fig. 2b). Our results suggest the possibility of photosynthetic competition for hydrogen sulphide between cyanobacteria and purple sulphur bacteria (PSB). Such competition could further explain why PSB thrive in autumn, whereas cyanobacteria and green non-sulphur bacteria (GNSB) are more abundant in summer. Interestingly, MAG 11 appears to be a phototroph capable of anoxygenic photosynthesis through hydrogen sulphide oxidation. Its genetic potential also suggests that it fixes nitrogen and carbon via the reductive pentose phosphate cycle, complementing the activities of cyanobacteria. This MAG does not belong to the PSB but is instead classified within the *Thiohalospiraceae* family (Supplementary Tables S8). Members of *Thiohalospiraceae* have been described exclusively as chemolithoautotrophs that obtain energy by oxidising inorganic sulphur compounds^54^. Thus, the discovery of photosynthesis-related genes in this MAG broadens the metabolic potential known for this family. Future isolation of an anoxygenic phototroph from *Thiohalospiraceae* would confirm this finding. MAGs 12 and 13, belonging to the order *Desulfobacterales*, are predicted to participate in both the reduction and oxidation parts of the sulphur cycle. MAG 13 also contains *acs* genes for carbon fixation via the reductive acetyl-CoA pathway, demonstrating that non-phototrophic groups also contribute to primary production.

The integration of the metagenomic analysis with the biogeochemical data greatly expanded our understanding of the Pozo Bravo microbial mats (Fig. 7). During the day, cyanobacteria and diatoms in the green surface layer perform oxygenic photosynthesis and assimilate inorganic carbon through the reductive pentose phosphate cycle. Certain cyanobacteria also fix atmospheric nitrogen and perform anoxygenic photosynthesis, releasing sulphate instead of oxygen. The oxygen generated in the green surface layer supports aerobic and facultative aerobic organisms, including chemoorganotrophic eukaryotes (Supplementary Table S10), which consume the organic exudates. Some of these organisms oxidise reduced compounds of hydrogen, sulphur, nitrogen, arsenic, and manganese. Anoxygenic phototrophs in the deeper reddish-pink layer contribute to primary production by assimilating inorganic carbon via the reductive pentose phosphate cycle (PSB), the rTCA cycle 1 (purple non-sulphur bacteria: PNSB), and the 3-hydroxypropionate bicycle (GNSB and PNSB). During anoxygenic photosynthesis, PSB use thiosulphate, hydrogen sulphide, or elemental sulphur as electron donors, whereas GNSB and PNSB use organic compounds or hydrogen. In the bottom dark brown layer of the mat (Fig. 2b), anaerobic heterotrophs oxidise the organic exudates using iron, manganese, cobalt, chromium, or other oxidised compounds as terminal electron acceptors, while fermenters use organic carbon as both electron donor and acceptor. Once night falls, photosynthesis ceases, and the residual oxygen is rapidly depleted by respiration. As a result, the anoxic zone expands, creating favourable conditions for anaerobic heterotrophs. Among them, sulphate reducing bacteria (SRB) reduce sulphate or other oxidised sulphur compounds, contributing to the regeneration of hydrogen sulphide levels. Certain SRB assimilate inorganic carbon via the acetyl-CoA pathway, further linking the carbon and sulphur cycles in the mat. The metabolic activities observed in the mat play a crucial role in shaping its mineralisation. The spatial and temporal separation of these metabolisms, as observed in the mat, creates localised differences in the saturation index that influence where and when carbonate precipitation occurs. Seasonal shifts in community composition also alter the balance of microbial metabolisms, leading to cyclical patterns of organomineralization. Therefore, the dynamic interplay between microbial processes and environmental conditions ultimately determines the lithification of the mat.

**Fig. 7:**
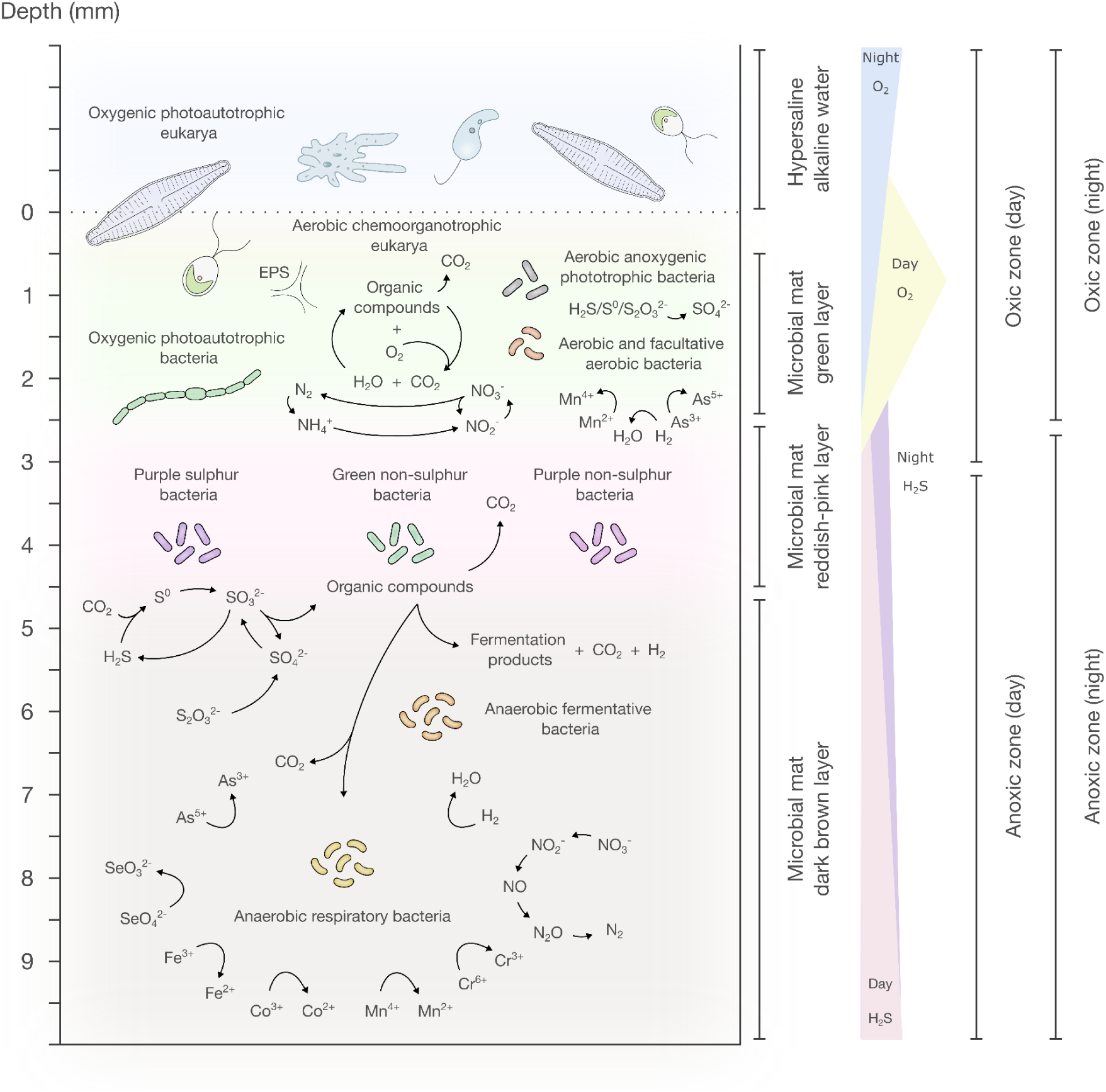
Schematic representation of Pozo Bravo soft microbial mat. The illustration depicts the major microbial guilds and biogeochemical cycles in the different layers of the microbial mat.

### Distinct prokaryotic compositions define modern microbialites around the world

To understand whether microbialites found in different locations around the world exhibit a uniform prokaryotic composition or whether their composition is influenced by geographic and environmental factors, we performed a global comparative analysis of modern microbialites from North America, South America, and Australia (Fig. 8). Our study reveals that these microbial structures do not share a similar prokaryotic composition; instead, their composition varies according to geographic location and environmental characteristics. For instance, the freshwater microbialites from Pavilion Lake^55^ contain 140 distinct prokaryotic genera characteristic of freshwater environments (Fig. 8b). Some members of these genera contribute to carbonate precipitation processes, including freshwater cyanobacteria (*Aphanocapsa*, *Chroococcus*, *Oscillatoria*, *Cyanomargarita*, and *Tolypothrix*) and sulphate reducers (*Sulfuricaulis*, *Sulfurisoma*, *Sulfurivirga*, *Desulforhabdus*, and *Bilophila*). The microbialites from Clifton Lake^56^, which develop under poikilosaline conditions, host 24 distinct prokaryotic genera, whose members are adapted to fluctuating salinity. These include halophilic or halotolerant cyanobacteria (*Gloeocapsa*, *Scytonema*, and *Chondrocystis*), anoxygenic phototrophs (*Chloracidobacterium*), and sulphate reducers (*Candidatus* Allobeggiato, *Desulfocella*, *Desulfospira*, and *Dissulfuribacter*) that drive carbonate precipitation. Microbialites from Pozo Bravo and Socompa^57^ lakes, which are exposed to similar conditions of hypersalinity, high radiation, low oxygen pressure, and slight alkalinity, share 8 distinct prokaryotic genera adapted to these extreme conditions (Fig. 8b). These genera include polyextremophilic anoxygenic phototrophs (*Thiorhodovibrio*) and sulphate reducers (*Dethiosulfatibacter*), which modify carbonate alkalinity and drive mineralization. Despite the shared extreme conditions, the microbialites from Pozo Bravo host 47 distinct prokaryotic genera not found in Socompa. These include polyextremophilic anoxygenic phototrophs (*Thiococcus*, *Thioflavicoccus*, and *Thiorhodococcus*) and sulphate reducers (*Pseudodesulfovibrio*) involved in carbonate precipitation, as well as genera involved in organic matter decomposition or fermentation (*Acetohalobium*, *Halanaerobaculum*, *Natronorubrum*, *Salinirubrum*, *Salimicrobium*, *Sporohalobacter*, *Anaerostipes*, *Faecalibacterium*, *Holdemanella*, *Intestinimonas*, *Succiniclasticum*, and *Agathobacter*). While the absence of these genera in Socompa reflects the influence of geographic location in shaping the prokaryotic community composition, interestingly, 14 prokaryotic genera are shared by all modern microbialites. These include genera involved in photosynthesis (*Thiohalocapsa* and *Candidatus* Alysiosphaera), sulphur reduction (*Desulfonema* and *Desulfovibrio*), and organic matter decomposition (*Sedimenticola*, *Sediminispirochaeta*, and *Peredibacter*), reflecting core ecological processes critical for microbialite formation and stability. The variability in prokaryotic compositions among modern microbialites demonstrates that the process of organomineralization depends not on the taxonomy of the organisms within the community but rather on their metabolic potential and the prevailing environmental conditions.

**Fig. 8:**
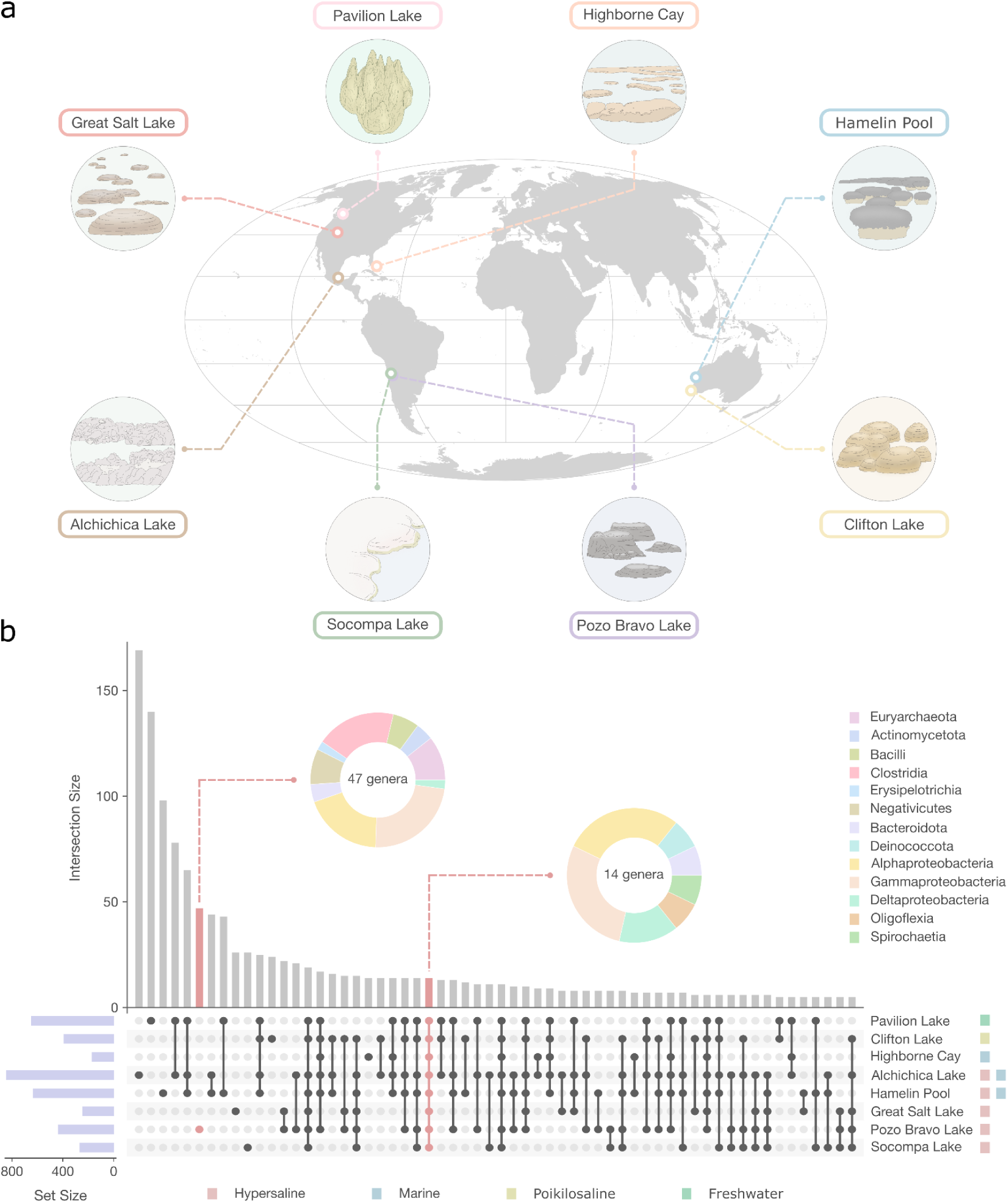
Modern microbialites around the world share some prokaryotic genera, but the degree of similarity varies depending on specific environmental conditions and locations. **a** Worldwide distribution of modern microbialites. **b** UpSet plot depicting common prokaryotic genera among modern microbialites around the globe. Vertical grey bars show the number of prokaryotic genera in each intersection, while horizontal purple bars display the number of prokaryotic genera in each set. The taxonomic classification of genera found only in Pozo Bravo’s microbialites and genera common to all microbialites is shown in the form of a pie chart.

## Conclusions

Pozo Bravo harbours a modern microbialite reef that develops under environmental conditions partially resembling those of the Precambrian era, making these microbialites one of the best models for the earliest biostructures on Earth. Therefore, the geobiological processes operating in this environment provide valuable insights into the mechanisms that shaped ancient microbialites. The formation of lithified mats and microbialites in Pozo Bravo is driven by microbial metabolic activities and environmental conditions that collectively determine their macro-, meso-, and microstructures, as well as their mineralogical composition. Ancient microbialites likely formed the same way, allowing us to use processes such as microbial precipitation of manganese oxides as potential biosignatures in the search for ancient life on Earth and the exploration of life beyond our planet. Microbial mats and microbialites in Pozo Bravo experience environmental fluctuations throughout the year, leading to seasonal variations in the microbial community composition. Both environmental changes and microbial successions influence organomineralization. In the warmer months, increased evaporation rates and higher abundances of photoautotrophs raise saturation indices, favouring carbonate precipitation. UV radiation also intensifies during this period, potentially destroying the EPS and lifting constraints on mineral precipitation. As a result, these seasonal dynamics create a cyclical pattern of organomineralization. Given that environmental cycles, including fluctuations in temperature, radiation levels, and evaporation rates, also existed in Precambrian Earth, it is highly probable that the earliest microbialites experienced similar periodic changes in microbial composition and organomineralization.

Although many geobiological processes that shape modern microbialites also played a fundamental role in the formation of ancient microbialites, some processes have changed in relevance over the course of Earth’s history. The metabolic repertoire of microbial communities, which is central to organomineralization, differed in ancient microbialites compared to most modern systems. Oxygenic photosynthesis carried out by cyanobacteria and unicellular eukaryotes, for instance, is a key driver of organomineralization in many modern microbialites. However, in the earliest microbialites, before the evolution of eukaryotic photoautotrophs, oxygenic photosynthesis may not have been the primary driver. Other processes, such as anoxygenic photosynthesis coupled to the oxidation of reduced sulphur or arsenic compounds, as observed in Pozo Bravo, could have been crucial alkaline engines in ancient Earth environments. Additionally, the presence of carbon fixation processes beyond the Calvin cycle in Pozo Bravo suggests that ancient microbialites may have assimilated inorganic carbon differently from most modern systems.

In summary, while many processes shaping ancient microbialites persist in modern systems, some key drivers of microbialite formation have evolved over time. Understanding both past and present geobiological processes is crucial for guiding our search for life beyond our planet and for interpreting the fossil record of Earth’s earliest biostructures.

## Methods

### Site description

In the southwest region of Laguna Pozo Bravo, microbial mats and microbialites are exposed to fluctuations in the water column, resulting in alternating moisture and drying periods. For this reason, soft (non-lithified) microbial mats, lithified microbial mats, and microbialites are interspersed, representing different stages of the lithification process (Fig. 1).

### Sample collection

Field campaigns to Laguna Pozo Bravo were carried out in May 2017 (fall), August 2017 (winter), November 2017 (spring), January 2018 (summer), January 2019 (summer), and January 2023 (summer) to collect samples and perform *in situ* analyses. The collected samples were selected based on a preliminary inspection of the sedimentary structures present throughout the lagoon. Water, soft microbial mat, and microbialite samples were collected in May 2017, August 2017, November 2017 and January 2018 for a seasonal 16S rRNA diversity analysis. Surface water and soft microbial mat samples were taken in August 2017 and January 2018 for chemical analyses. Additional soft microbial mat samples were collected in January 2023 for µXRF, SEM, metagenomic, and bulk geochemistry analyses. Lithified microbial mats and microbialites were sampled in January 2019 and January 2023 for mineralogy and Raman spectroscopy analyses, respectively.

Samples for 16S rRNA diversity and metagenomic analyses were collected employing autoclaved stainless steel tweezers and spoon spatulas, and transferred to sterile 50 ml tubes or polyethylene containers. These samples were stored in RNAlater® solution (Thermo Fisher Scientific, United States) at 4°C in the dark and processed within a week. Samples for µXRF, SEM, mineralogy, Raman spectroscopy, chemical, and bulk geochemistry analyses were collected employing solvent-cleaned (dichloromethane and methanol) stainless steel tweezers and spoon spatulas, and transferred to polyethylene containers. These samples were kept at 4°C in the dark until analysis in the laboratory.

### Physicochemical analysis

Environmental parameters (pH, conductivity, temperature, and dissolved oxygen) of both surface water and soft microbial mats were measured *in situ* in August 2017 and January 2018 using a portable Hanna HI 9829 multiparameter probe. The instrument was calibrated according to the manufacturer’s instructions prior to measurement. Water-level fluctuations along the shoreline were monitored by placing PVC pipes at the lagoon banks and measuring the water column height. Chemical parameters of the water and soft microbial mats were analysed by Grupo Induser S.R.L (Salta, Argentina), and the corresponding values are presented in Tables 1 and 2. Using the physicochemical parameters of the water, mineral saturation indices were modelled with PHREEQC v3.0 geochemical software^58^. The saturation index (SI) is defined as SI=log(IAP/K_sp_), where IAP represents the ion activity product, and K_sp_ the solubility product constant of the corresponding mineral. If SI<0 the solution is undersaturated with respect to the mineral, if SI=0 the solution is at equilibrium with the mineral, and if SI>0 the solution is supersaturated with respect to the mineral.

### Satellite data acquisition and analysis

Surface pressure (monthly) [v5.12.4 STD], UV index (local noon) [standard v003], and land surface temperature (L3, monthly, day/night, TES algorithm) [v6.1 STD] data for the Laguna Pozo Bravo region were obtained from NASA Earthdata (https://earthdata.nasa.gov/). Data processing was conducted using the R packages *rhdf5* and *ncdf4*. The *rhdf5* package was used to extract data from HDF5 format files, while the *ncdf4* package facilitated handling netCDF format files. Data visualisation was carried out using the R package *ggplot2*.

### Microelectrode measurements

Oxygen and sulphide profiles covering the upper 5–10 mm of the soft microbial mats were measured *in situ* in triplicate during both daytime (from 13:00 to 16:00) and nighttime (from 20:00 to 01:00) in January 2019. Clark-type oxygen and amperometric sulphide microsensors with a tip width of <50 µm were employed for these measurements, constructed, calibrated, and operated as previously described^59–62^. The microsensors were placed on a multisensor holder with a tip distance of ∼0.5 cm for simultaneous measurements, with electricity supplied by a portable generator. Oxygen concentration was adjusted for salinity and temperature, according to specifications (Unisense), and values were determined assuming an atmospheric pressure of 1 bar^63^.

### Scanning micro X-ray fluorescence analysis

The distribution of elements within a freeze-dried soft microbial mat sample was determined by micro X-ray fluorescence spectrometry employing an M4 Tornado μXRF spectrometer (Bruker Nano GmbH, Germany). The instrument was equipped with a Rh X-ray source and polycapillary optics (20 µm spot size). Scanning was operated at 50 kV and 600 µA tube setting under vacuum condition of 20 mb. The pixel size was set to 25 µm, and scan time was 20 ms/pix. Elemental distributions were analysed as net intensities (deconvoluted counts) using the Bruker M4 Esprit software package.

### Raman Spectroscopy

Raman spectroscopy analysis was performed on a microbialite sample to assess mineralogical variations through both macro- and microscopic observations. Spectra were acquired using backscattering geometry with a LabRam HR Evolution Raman microspectrometer, set to a resolution of 0.4 cm^−1^ and equipped with a He-Ne laser line (633 nm) at INQUIMAE (UBA-CONICET), Buenos Aires, Argentina. Beam power and acquisition times were optimised for each sample to obtain informative spectra without causing sample alteration. Spectra were recorded using a 50× microscope objective (spatial resolution of approximately 1 μm), with an exposure time of 10-30 s and three accumulations.

### Mineralogy and bulk geochemistry analyses

The mineral composition of freeze-dried (soft and lithified) microbial mat and microbialite samples was determined through powder X-ray diffraction (XRD) analysis using a Bruker X-ray diffractometer (Eco D8 Advance, XRD), equipped with a Cu X-ray source (Cu Kα1,2, λ=1.54056 Å), operated at 40 kV and 25 mA. The freeze-dried and finely ground (<20 μm) samples were scanned from 5° to 60° in the 2·ϴ-diffraction angle, with a scanning step size of 0.02° and a time per step of 1 s. Diffraction patterns were identified using the DIFFRAC-EVA software, with reference to the PDT-2 2002 database included in the software. Semi-quantification of the minerals was performed based on the intensity of the main peaks in each XRD pattern^64,65^. The mineralogy analysis of the microbialite sample was further complemented with microscopic observations using conventional optical microscopy.

The stable isotopic composition of organic carbon (δ^13^C) and total nitrogen (δ^15^N) in bulk soft microbial mat and microbialite samples was determined using isotope-ratio mass spectrometry (IRMS), following the method outlined by the United States Geological Survey (USGS)^66^. Samples were first homogenised by manual grinding in a mortar and then decarbonated with HCl (3N). After 24 hours of equilibration, the samples were adjusted to neutral pH using ultrapure water and subsequently dried in an oven at 50 °C until a constant weight was achieved. The δ^13^C and δ^15^N ratios were measured using a MAT 253 IRMS (Thermo Fisher Scientific, Waltham, Massachusetts, USA) and reported in standard per mil notation (‰) relative to international reference standards (δ^13^C vs. Vienna Pee Dee Belemnite and δ^15^N vs. air). Three certified standards (USGS41, IAEA-600, and USGS40) were employed with an analytical precision of 0.1‰. Total organic carbon (TOC %) and total nitrogen (TN %) content were determined using an elemental analyser (HT Flash, Thermo Fisher Scientific, Waltham, Massachusetts, USA) during the stable isotope measurements. The mineralogical and isotopic analysis have been done at the Centro de Astrobiología (CAB), in Madrid (Spain).

### Scanning electron microscopy analysis

SEM images of the soft microbial mats were obtained under vacuum conditions using a Zeiss Supra 55 VP scanning electron microscope (Carl Zeiss NTS GmbH, Germany) at Centro Integral de Microscopía Electrónica (CIME) in Tucumán, Argentina. Prior to observation, samples were fixed overnight at 4°C in modified Karnovsky fixative, composed of 8% (v/v) formaldehyde, 16% (v/v) glutaraldehyde, and phosphate-buffered saline (200 mM PBS, pH 7.4). Samples were then washed three times with PBS and calcium chloride (CaCl_2_) for 10 min each, followed by fixation with 2% (v/v) osmium tetroxide overnight. After two ethanol washes (30% v/v) for 10 min each, the samples were dried at the critical point and sputtered with gold.

Diatom taxa were identified using SEM images and identification keys^67^, based on morphological characteristics of their frustules. The analysed morphological features included valve size, shape, and symmetry; the presence, location and structure of the axial area; the presence, location, and structure of the raphe; the presence of septa; the presence and structure of costae; the location, structure, and density of striae; and the presence of stauros, among others.

### DNA extraction and sequencing

Different procedures were employed for DNA extraction depending on the type of sample. Water samples were filtered through a 0.22-µm pore size membrane (DURAPORE GV) to concentrate suspended microorganisms. Soft microbial mat samples were homogenised to ensure representativeness. While microbialite samples were immersed in phosphate-buffered saline (50 mM PBS, pH 7.5) and sonicated three times to detach microorganisms from the surface. The supernatants were then collected and centrifuged at 13,000 g for 15 min. The resulting cell pellets served as starting material for DNA extraction.

Total genomic DNA was isolated using the FastDNA® SPIN Kit for Soil (MP Biomedicals, United States) following the manufacturer’s protocol. DNA quality and concentration were assessed using a NanoDrop 2000c Spectrophotometer (Thermo Fisher Scientific, United States). The hypervariable V3 and V4 regions of the bacterial and archaeal 16S rRNA gene were amplified using the primers Bakt_341F (5-CCTACGGGNGGCWGCAG) and Bakt_805R (5-GACTACHVGGGTATCTAATCC). Amplicons obtained from water, soft microbial mat and microbialite samples, along with total DNA isolated from a soft microbial mat sample, were sequenced using an Illumina MiSeq platform. Raw sequences were deposited in the ENA Project database under the accession numbers ERP120954 (amplicon sequences) and ERP113390 (metagenomic sequences).

### Amplicon sequence analysis

The prokaryotic composition of water, soft microbial mat and microbialite samples (ENA accession number: ERP120954) was determined with the MGnify metagenomic analysis pipeline v.5.0^68^ (MGnify accession number: MGYS00005312) through the taxonomic assignment of the 16S rRNA gene amplicon sequences (SILVA release 132)^69^. Alpha diversity metrics (Shannon index)^70^ were calculated using the R package *phyloseq*^71^. The *phyloseq* package was also employed to perform a principal coordinate analysis (PCoA) based on Bray-Curtis dissimilarity^72^. Permutational ANOVA (PERMANOVA) tests were carried out using the R package *vegan*^73^ to evaluate the PCoA result. To compare the genera of prokaryotes found in Pozo Bravo’s microbialites with those in modern microbialites from other locations around the world - Alchichica Lake, Puebla, Mexico (SRP061655, SRP072547)^74–76^; Great Salt Lake, Utah, United States (SRP067068)^77^; Highborne Cay, Bahamas (SRP004035)^78^; Pavilion Lake, British Columbia, Canada (SRP035880, ERP020021)^55^; Hamelin Pool, Shark Bay, Australia (SRP055055)^79^; Clifton Lake, Yalgorup National Park, Australia (SRP072185)^56^; and Socompa Lake, Salta, Argentina (SRP007748, SRP072938, ERP021393)^57,80^ - an UpSet plot was generated using the R package *UpSetR*^81^.

### Metagenomic analysis

Quality control checks on the raw paired-end sequence data (ENA accession number: ERP113390) were first conducted using FastQC v.0.11.9 (Babraham Bioinformatics website). The results from this analysis were later used to determine the necessary trimming steps (ILLUMINACLIP = TruSeq2-PE.fa:2:30:5, SLIDINGWINDOW = 4:20, AVGQUAL = 20, and MINLEN = 50), which were carried out with Trimmomatic v.0.39^82^. Taxonomic and functional annotations of the cleaned paired reads were performed using the MGnify metagenomic analysis pipeline v.4.1^68^ (MGnify accession number: MGYS00004549). The cleaned paired reads were further assembled into contigs using MetaSPAdes v.3.14.1^83^, and the quality of the assembly (Supplementary Table S6) was assessed with QUAST v.4.6^84^ and Bowtie2 v.2.4.2^85^. Metagenome-assembled genomes (MAGs) were reconstructed from the contigs employing MetaWRAP v.1.1^86^. Quality control checks on the MAGs were conducted using CheckM v.1.0.13^87^ (Supplementary Table S7). Using the quality control metrics, dereplication of MAGs was conducted with dRep v.1.4.3^88^. This tool identifies groups of MAGs with high similarity (ANI, Average Nucleotide Identity > 95%) and selects the representative MAG from each group. In this way, redundant MAGs are filtered out. Taxonomic assignment of MAGs was performed by comparison against the Genome Database Taxonomy (GTDB) release 220^89^ (Supplementary Table S8). The prediction and functional annotation of protein-coding genes in the assembled metagenome and MAGs were performed with Prodigal v.2.6.3^90^ and InterProScan v.5.34-73.0^91^, respectively. After the functional annotation of the genes, an analysis of the metabolic pathways present in the MAGs was carried out with Genome Properties v.2.0^92^. Genes involved in metabolic pathways/cycles of interest were additionally annotated with HMMER-3.1 (http://hmmer.org/) using hidden Markov model (HMM) profiles for KEGG/KO with predefined score thresholds^93^ (Supplementary Table S9). Cleaned paired reads were mapped to the identified genes using Bowtie2 v.2.4.2^85^, and the total number of mapped reads obtained was used to determine the abundance of each gene, normalised by gene length and library size [Reads per kilobase per million mapped reads (RPKM)].

## Acknowledgements

We acknowledge the support of the EMBL Planetary Biology Transversal Theme through the seed grant awarded to F.A.V., L.S-G., and M.M.G-A., as well as the CABANA project, funded by UKRI-BBSRC on behalf of the Global Challenges Research Fund (BB/P027849/1). We thank Luis Ahumada, Agustina I. Lencina, Mariana N. Soria, and the native communities of Antofalla, El Peñón, and Antofagasta de la Sierra for their support during field work. We also appreciate María Ana Castro for her assistance with Raman analysis. We acknowledge the technical support from the Centro Integral de Microscopía Electrónica (CIME) in Tucumán, Argentina, for SEM experiments and the SPC facility at EMBL Hamburg for additional technical assistance. We express our gratitude to the “Secretaría de Medio Ambiente de la Provincia de Catamarca” for their administrative support that allowed our sampling (Expte.EX-2022-02222431-CAT-DPB#SEAS, Resolución S.E.A., D.S. N°: 053/2017). Additionally, we thank the “Secretaría de Política Ambiental en Recursos Naturales, Ministerio de Ambiente y Desarrollo Sostenible de la Nación Argentina” for providing the certificate of compliance (Title: IF-2023-65966642-APN-SPARN#MAD, UId: ABSCH-IRCC-AR-264943-1) and the export certificate for genetic resources (CE-2023-69835755-APN-SPARN#MAD).

## Author Contributions

F.A.V., M.M.G-A., and M.E.F. designed the project and coordinated the research activities. F.A.V. and M.M.G-A. wrote the manuscript with input from all co-authors. F.A.V. was involved in all data analyses of the study. F.A.V., L.S-G., D.C., A.C.S., J.M.K., and M.E.F. collected the samples. A.C.S. and J.M.K. conducted the microelectrode measurements. A.C.S., H.T., and J.M.K. performed the µXRF analysis. S.O. carried out the Raman spectroscopy analysis. L.S-G. and D.C. conducted the mineralogy and bulk geochemistry analyses. F.A.V. extracted the total DNA from the samples. F.A.V., A.L.M., and R.D.F. performed the bioinformatic analyses. L.S-G., R.D.F., A.G.T., M.M.G-A., and M.E.F. obtained funding to carry out the project.

## References

1. Microbial Ecosystems in Central Andes Extreme Environments: Biofilms, Microbial Mats, Microbialites and Endoevaporites. (Springer, Cham, 2020).

2. Lyons, T. W. et al. Co-evolution of early Earth environments and microbial life. Nat Rev Microbiol 22, 572–586 (2024).

3. Sauterey, B., Charnay, B., Affholder, A., Mazevet, S. & Ferrière, R. Co-evolution of primitive methane-cycling ecosystems and early Earth’s atmosphere and climate. Nat Commun 11, 2705 (2020).

4. Cabrol, N. The evolution of lacustrine environments on mars: Is mars only hydrologically dormant? Icarus 149, 291–328 (2001).

5. Wharton, R. A., Crosby, J. M., McKay, C. P. & Rice, J. W. Paleolakes on Mars. Journal of Paleolimnology vol. 13 267–283 Preprint at 10.1007/bf00682769 (1995).

6. Sánchez-García, L. et al. Assessing siliceous sinter matrices for long-term preservation of lipid biomarkers in opaline sinter deposits analogous to Mars in El Tatio (Chile). Sci. Total Environ. 870, 161765 (2023).

7. Squyres, S. W. et al. Detection of silica-rich deposits on Mars. Science 320, 1063–1067 (2008).

8. Carrizo, D., Vignale, F. A., Sánchez-García, L. & Farías, M. E. Ecological variability based on lipid biomarkers in astrobiologically interesting wetlands from the Argentinian central Andes. FEMS Microbiol. Ecol. 98, (2022).

9. Vignale, F. A. et al. Lithifying and Non-Lithifying Microbial Ecosystems in the Wetlands and Salt Flats of the Central Andes. Microb. Ecol. (2021) doi:10.1007/s00248-021-01725-8.

10. López, D., Vlamakis, H. & Kolter, R. Biofilms. Cold Spring Harb. Perspect. Biol. 2, a000398 (2010).

11. Jain, A., Gupta, Y., Agrawal, R., Khare, P. & Jain, S. K. Biofilms--a microbial life perspective: a critical review. Crit. Rev. Ther. Drug Carrier Syst. 24, 393–443 (2007).

12. Prieto-Barajas, C. M., Valencia-Cantero, E. & Santoyo, G. Microbial mat ecosystems: Structure types, functional diversity, and biotechnological application. Electron. J. Biotechnol. 31, 48–56 (2018).

13. Madigan, M. T., Aiyer, J., Buckley, D. H., Matthew Sattley, W. & Stahl, D. A. Brock Biology of Microorganisms, Global Edition. (Pearson Higher Ed, 2021).

14. Noffke, N., Christian, D., Wacey, D. & Hazen, R. M. Microbially induced sedimentary structures recording an ancient ecosystem in the ca. 3.48 billion-year-old Dresser Formation, Pilbara, Western Australia. Astrobiology 13, 1103–1124 (2013).

15. Buongiorno, J., Gomez, F. J., Fike, D. A. & Kah, L. C. Mineralized microbialites as archives of environmental evolution, Laguna Negra, Catamarca Province, Argentina. Geobiology 17, 199–222 (2019).

16. Burne, R. V. & Moore, L. S. Microbialites; organosedimentary deposits of benthic microbial communities. Palaios 2, 241–254 (1987).

17. Dupraz, C. & Visscher, P. T. Microbial lithification in marine stromatolites and hypersaline mats. Trends Microbiol. 13, 429–438 (2005).

18. Dupraz, C. et al. Processes of carbonate precipitation in modern microbial mats. Earth-Sci. Rev. 96, 141–162 (2009).

19. Microbial Sediments. (Springer, Berlin, Heidelberg, 2000).

20. Shapiro, R. S. A comment on the systematic confusion of thrombolites. Palaios 15, 166 (2000).

21. Suosaari, E. P., Awramik, S. M., Reid, R. P., Stolz, J. F. & Grey, K. Living dendrolitic microbial mats in Hamelin pool, shark bay, western Australia. Geosciences (Basel) 8, 212 (2018).

22. Braga, J. C., Martin, J. M. & Riding, R. Controls on microbial dome fabric development along a carbonate-siliciclastic shelf-basin transect, Miocene, SE Spain. Palaios 10, 347 (1995).

23. Grey, K. & Awramik, S. M. Handbook for the Study and Description of Microbialites. (Geological Survey of Western Australia, Bulletin 147, 2020).

24. Riding, R. Microbial carbonates: the geological record of calcified bacterial–algal mats and biofilms. Sedimentology 47, 179–214 (2000).

25. Riding, R. Classification of Microbial Carbonates. in Calcareous Algae and Stromatolites (ed. Riding, R.) 21–51 (Springer Berlin Heidelberg, Berlin, Heidelberg, 1991).

26. Feldmann, M. & McKenzie, J. A. Stromatolite-thrombolite associations in a modern environment, lee stocking island, Bahamas. Palaios 13, 201 (1998).

27. Moore, L. S. & Burne, R. V. The modern thrombolites of lake Clifton, western Australia. in Phanerozoic Stromatolites II 3–29 (Springer Netherlands, Dordrecht, 1994).

28. Riding, R. Microbial carbonate abundance compared with fluctuations in metazoan diversity over geological time. Sediment. Geol. 185, 229–238 (2006).

29. Walter, M. R. & Heys, G. R. Links between the rise of the metazoa and the decline of stromatolites. Precambrian Res. 29, 149–174 (1985).

30. Monty, C. Precambrian background and Phanerozoic history of stromatolitic communities, an overview. asgb (1974).

31. Awramik, S. M. The history and significance of stromatolites. in Early Organic Evolution 435–449 (Springer Berlin Heidelberg, Berlin, Heidelberg, 1992).

32. Reid, R. P., Suosaari, E. P., Oehlert, A. M., Pollier, C. G. L. & Dupraz, C. Microbialite Accretion and Growth: Lessons from Shark Bay and the Bahamas. Ann Rev Mar Sci 16, 487–511 (2024).

33. Lopes, F., Courtillot, V. & Le Mouël, J.-L. Triskeles and symmetries of mean global sea-level pressure. Atmosphere (Basel) 13, 1354 (2022).

34. Luo, G. et al. Rapid oxygenation of Earth’s atmosphere 2.33 billion years ago. Sci Adv 2, e1600134 (2016).

35. Cockell, C. S. The ultraviolet history of the terrestrial planets — implications for biological evolution. Planet. Space Sci. 48, 203–214 (2000).

36. Munguira, A. et al. Near surface atmospheric temperatures at Jezero from Mars 2020 MEDA measurements. J. Geophys. Res. Planets 128, (2023).

37. Knauth, L. P. Temperature and salinity history of the Precambrian ocean: implications for the course of microbial evolution. in Geobiology: Objectives, Concepts, Perspectives 53–69 (Elsevier, 2005).

38. Grotzinger, J. P. & Kasting, J. F. New constraints on Precambrian ocean composition. J Geol 101, 235–243 (1993).

39. Sforna, M. C. et al. Evidence for arsenic metabolism and cycling by microorganisms 2.7 billion years ago. Nat. Geosci. 7, 811–815 (2014).

40. Tostevin, R. & Ahmed, I. A. M. Micronutrient availability in Precambrian oceans controlled by greenalite formation. Nat. Geosci. (2023) doi:10.1038/s41561-023-01294-0.

41. Ehlmann, B. L. & Edwards, C. S. Mineralogy of the martian surface. Annu. Rev. Earth Planet. Sci. 42, 291–315 (2014).

42. Lencina, A. I., Soria, M. N., Gomez, F. J., Gérard, E. & Farias, M. E. Composite microbialites: Thrombolite, dendrolite, and stromatolite associations in a modern environment, Pozo Bravo lake, Salar de Antofalla, Catamarca Puna, Argentina. J. Sediment. Res. 91, 1305–1330 (2021).

43. Sujith, P. P. & Bharathi, P. A. L. Manganese oxidation by bacteria: biogeochemical aspects. Prog Mol Subcell Biol 52, 49–76 (2011).

44. Sjöberg, S. et al. Microbe-mediated Mn oxidation—A proposed model of mineral formation. Minerals (Basel) 11, 1146 (2021).

45. Mitra, K., Moreland, E. L., Ledingham, G. J. & Catalano, J. G. Formation of manganese oxides on early Mars due to active halogen cycling. Nat. Geosci. 16, 133–139 (2023).

46. Gasda, P. J. et al. Manganese-rich sandstones as an indicator of ancient oxic lake water conditions in Gale crater, Mars. J. Geophys. Res. Planets 129, (2024).

47. Thomassin, J. H. & Touray, J. C. Hydrotalcite, a temporary hydroxycarbonate early developed during the interaction basaltic glass sea-water. Bulletin De Mineralogie 105, 312–319 (1982).

48. Calder, J. A. & Parker, P. L. Geochemical implications of induced changes in C13 fractionation by blue-green algae. Geochim. Cosmochim. Acta 37, 133–140 (1973).

49. Pardue, J. W., Scalan, R. S., Van Baalen, C. & Parker, P. L. Maximum carbon isotope fractionation in photosynthesis by blue-green algae and a green alga. Geochim. Cosmochim. Acta 40, 309–312 (1976).

50. Thompson, P. A. & Calvert, S. E. Carbon-isotope fractionation by a marine diatom: The influence of irradiance, daylength, pH, and nitrogen source. Limnol. Oceanogr. 39, 1835–1844 (1994).

51. Hayes, J. M. Fractionation of Carbon and Hydrogen Isotopes in Biosynthetic Processes. Reviews in Mineralogy and Geochemistry vol. 43 225–277 Preprint at 10.2138/gsrmg.43.1.225 (2001).

52. Robinson, D. delta(15)N as an integrator of the nitrogen cycle. Trends Ecol. Evol. 16, 153–162 (2001).

53. Beblawy, S. et al. Extracellular reduction of solid electron acceptors by Shewanella oneidensis. Mol Microbiol 109, 571–583 (2018).

54. Sorokin, D. Y. & Merkel, A. Y. *Thiohalospiraceae fam. nov. Bergey’s Manual of Systematics of Archaea and Bacteria* 1–2 Preprint at 10.1002/9781118960608.fbm00397 (2023).

55. Schulze-Makuch, D. et al. Pavilion lake microbialites: morphological, molecular and biochemical evidence for a cold-water transition to colonial aggregates. Life (Basel) 3, 21–37 (2012).

56. Warden, J. G. et al. Characterization of Microbial Mat Microbiomes in the Modern Thrombolite Ecosystem of Lake Clifton, Western Australia Using Shotgun Metagenomics. Front Microbiol 7, 1064 (2016).

57. Farías, M. E. et al. The discovery of stromatolites developing at 3570 m above sea level in a high-altitude volcanic lake Socompa, Argentinean Andes. PLoS One 8, e53497 (2013).

58. Parkhurst, D. L. & Appelo, C. A. J. USGS - Description of Input and Examples for PHREEQC Version 3—A Computer Program for Speciation, Batch-Reaction, One-Dimensional Transport, and Inverse Geochemical Calculations. U.S. Geological Survey (USGS)Description of Input and Examples for PHREEQC Version 3—A Computer Program for Speciation, Batch-Reaction, One-Dimensional Transport, and Inverse Geochemical Calculations https://pubs.usgs.gov/tm/06/a43/ (2013).

59. De Beer, D., Schramm, A., Santegoeds, C. M. & Kuhl, M. A nitrite microsensor for profiling environmental biofilms. Appl. Environ. Microbiol. 63, 973–977 (1997).

60. Jeroschewski, P., Steuckart, C. & Kühl, M. An amperometric microsensor for the determination of H2S in aquatic environments. Anal. Chem. 68, 4351–4357 (1996).

61. Klatt, J. M. et al. Structure and function of natural sulphide-oxidizing microbial mats under dynamic input of light and chemical energy. ISME J. 10, 921–933 (2016).

62. Revsbech, N. P. An oxygen microsensor with a guard cathode. Limnol. Oceanogr. 34, 474–478 (1989).

63. Forstner, H. & Gnaiger, E. Calculation of equilibrium oxygen concentration. in Polarographic Oxygen Sensors 321–333 (Springer Berlin Heidelberg, Berlin, Heidelberg, 1983).

64. Schultz, L. G. Quantitative interpretation of mineralogical composition from X-ray and chemical data for the Pierre Shale. U.S. Geol. Surv. Prof. Pap. 391–C (1964).

65. Moore, D. M., Reynolds, R. C., Jr & Others. X-Ray Diffraction and the Identification and Analysis of Clay Minerals. (Oxford University Press (OUP), 1989).

66. Revesz, K., Qi, H. & Coplan, T. B. Determination of the δ^15^N and δ^13^C of total nitrogen and carbon in solids; RSIL lab code 1832. Techniques and Methods Preprint at 10.3133/tm10c5 (2006).

67. Modern Trends in Diatom Identification. (Springer, Cham, 2020).

68. Mitchell, A. L. et al. MGnify: the microbiome analysis resource in 2020. Nucleic Acids Res. 48, D570–D578 (2020).

69. Glöckner, F. O. et al. 25 years of serving the community with ribosomal RNA gene reference databases and tools. J Biotechnol 261, 169–176 (2017).

70. Shannon, C. E. A mathematical theory of communication. Bell Syst. Tech. J. 27, 379–423 (1948).

71. McMurdie, P. J. & Holmes, S. phyloseq: an R package for reproducible interactive analysis and graphics of microbiome census data. PLoS One 8, e61217 (2013).

72. Bray, J. R. & Curtis, J. T. An ordination of the upland forest communities of southern Wisconsin. Ecol. Monogr. 27, 325–349 (1957).

73. Dixon, P. VEGAN, a package of R functions for community ecology. J. Veg. Sci. 14, 927–930 (2003).

74. Saghaï, A. et al. Metagenome-based diversity analyses suggest a significant contribution of non-cyanobacterial lineages to carbonate precipitation in modern microbialites. Front Microbiol 6, 797 (2015).

75. Saghaï, A., Zivanovic, Y., Moreira, D., Tavera, R. & López-García, P. A Novel Microbialite-Associated Phototrophic Chloroflexi Lineage Exhibiting a Quasi-Clonal Pattern along Depth. Genome Biol Evol 12, 1207–1216 (2020).

76. Saghaï, A. et al. Comparative metagenomics unveils functions and genome features of microbialite-associated communities along a depth gradient. Environ Microbiol 18, 4990–5004 (2016).

77. Lindsay, M. R., Dunham, E. C. & Boyd, E. S. Microbialites of great Salt Lake. in Great Salt Lake Biology 87–118 (Springer International Publishing, Cham, 2020).

78. Baumgartner, L. K. et al. Microbial diversity in modern marine stromatolites, Highborne Cay, Bahamas. Environmental Microbiology 11, 2710–2719 (2009).

79. Suosaari, E. P. et al. New multi-scale perspectives on the stromatolites of Shark Bay, Western Australia. Sci Rep 6, 20557 (2016).

80. Toneatti, D. M., Albarracín, V. H., Flores, M. R., Polerecky, L. & Farías, M. E. Stratified Bacterial Diversity along Physico-chemical Gradients in High-Altitude Modern Stromatolites. Front. Microbiol. 8, 646 (2017).

81. Conway, J. R., Lex, A. & Gehlenborg, N. UpSetR: an R package for the visualization of intersecting sets and their properties. Bioinformatics 33, 2938–2940 (2017).

82. Bolger, A. M., Lohse, M. & Usadel, B. Trimmomatic: a flexible trimmer for Illumina sequence data. Bioinformatics 30, 2114–2120 (2014).

83. Bankevich, A. et al. SPAdes: a new genome assembly algorithm and its applications to single-cell sequencing. J. Comput. Biol. 19, 455–477 (2012).

84. Gurevich, A., Saveliev, V., Vyahhi, N. & Tesler, G. QUAST: quality assessment tool for genome assemblies. Bioinformatics 29, 1072–1075 (2013).

85. Langmead, B. & Salzberg, S. L. Fast gapped-read alignment with Bowtie 2. Nat. Methods 9, 357–359 (2012).

86. Uritskiy, G. V., DiRuggiero, J. & Taylor, J. MetaWRAP-a flexible pipeline for genome-resolved metagenomic data analysis. Microbiome 6, 158 (2018).

87. Parks, D. H., Imelfort, M., Skennerton, C. T., Hugenholtz, P. & Tyson, G. W. CheckM: assessing the quality of microbial genomes recovered from isolates, single cells, and metagenomes. Genome Res. 25, 1043–1055 (2015).

88. Olm, M. R., Brown, C. T., Brooks, B. & Banfield, J. F. dRep: a tool for fast and accurate genomic comparisons that enables improved genome recovery from metagenomes through de-replication. ISME J. 11, 2864–2868 (2017).

89. Parks, D. H. et al. GTDB: an ongoing census of bacterial and archaeal diversity through a phylogenetically consistent, rank normalized and complete genome-based taxonomy. Nucleic Acids Res. 50, D785–D794 (2022).

90. Hyatt, D. et al. Prodigal: prokaryotic gene recognition and translation initiation site identification. BMC Bioinformatics 11, 119 (2010).

91. Jones, P. et al. InterProScan 5: genome-scale protein function classification. Bioinformatics 30, 1236–1240 (2014).

92. Richardson, L. J. et al. Genome properties in 2019: a new companion database to InterPro for the inference of complete functional attributes. Nucleic Acids Res. 47, D564–D572 (2019).

93. Aramaki, T. et al. KofamKOALA: KEGG Ortholog assignment based on profile HMM and adaptive score threshold. Bioinformatics 36, 2251–2252 (2020).

