## Supplementary information for "Seasonal biogeochemical variations in a modern microbialite reef under early Earth-like conditions"

**Supplementary Figures**

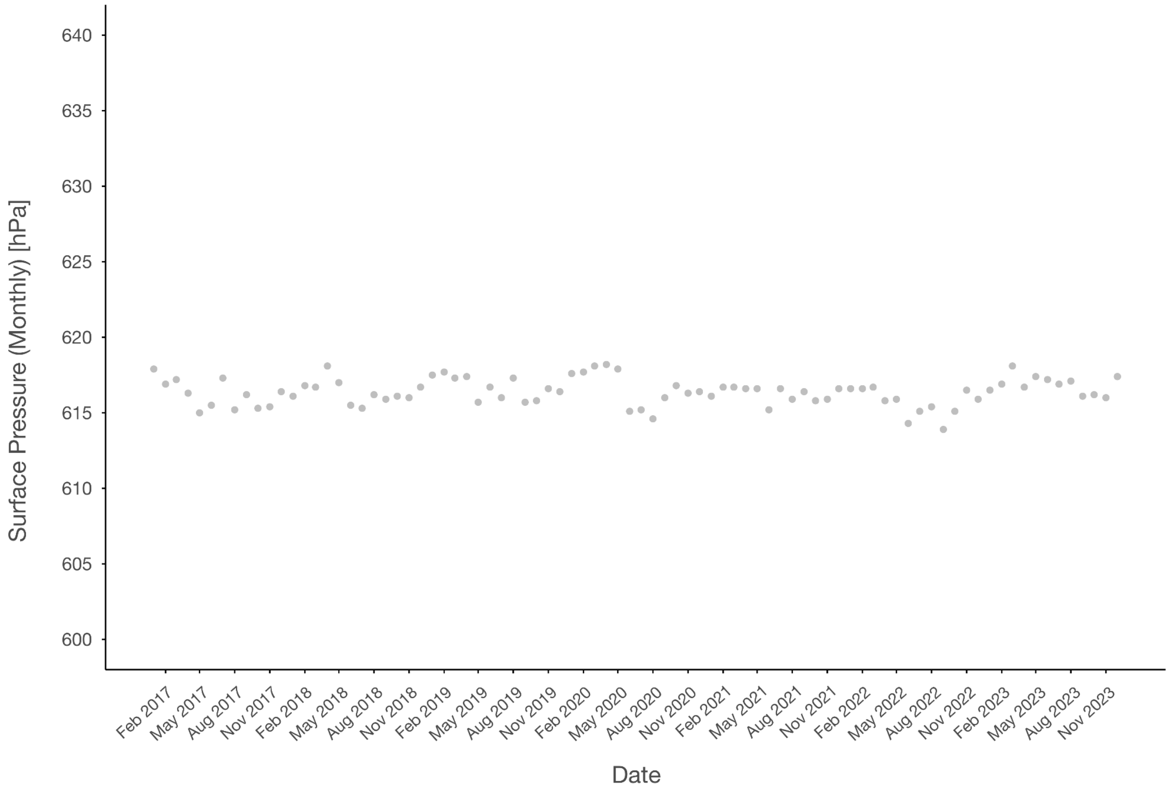


**Supplementary Figure S1. Monthly surface pressure in the Laguna Pozo Bravo region is consistently low (~60% of sea level pressure) throughout the seasons.** Monthly surface-level atmospheric pressure in hectopascals (hPa) for the Laguna Pozo Bravo region from January 2017 to December 2023.

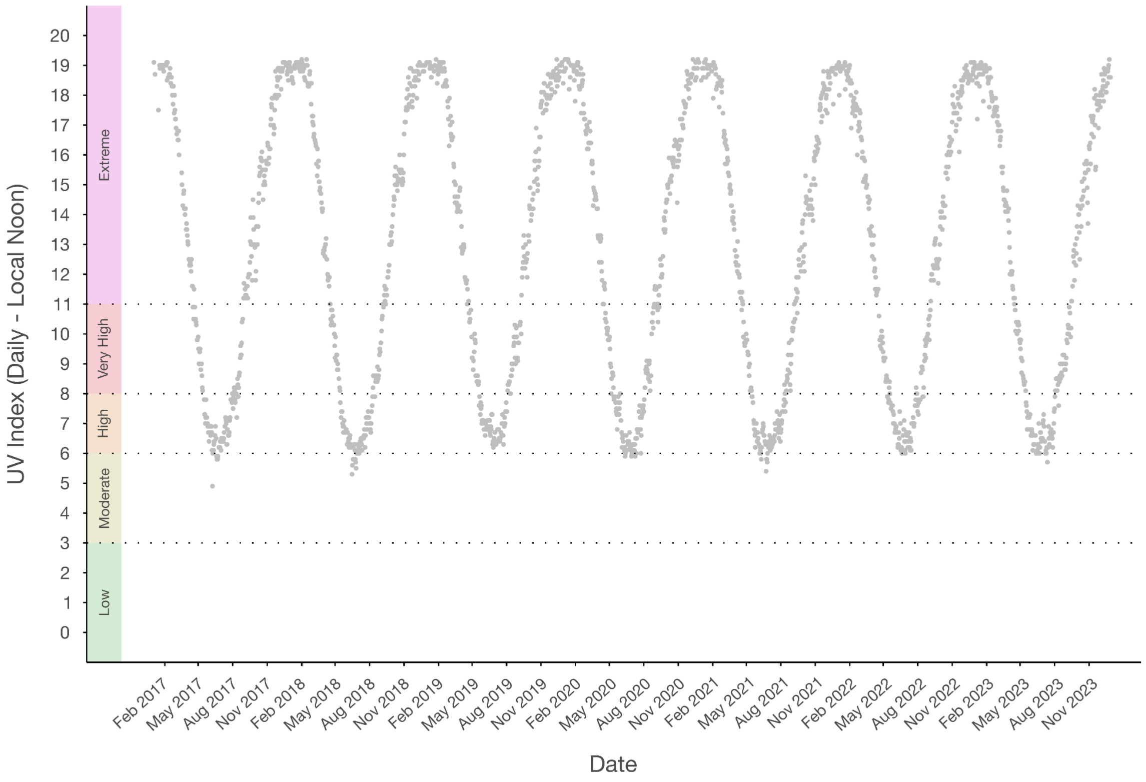


**Supplementary Figure S2. The daily UV index in the Laguna Pozo Bravo region varies between seasons, showing high values in winter and extreme values in summer.** Daily UV index (local noon) for the Laguna Pozo Bravo region from January 2017 to December 2023. Low (green), Medium (yellow), High (light orange), Very High (dark orange) and Extreme (pink) intervals are defined based on its potential risk to human health (WHO: Global Solar UV Index: A Practical Guide, 2002).

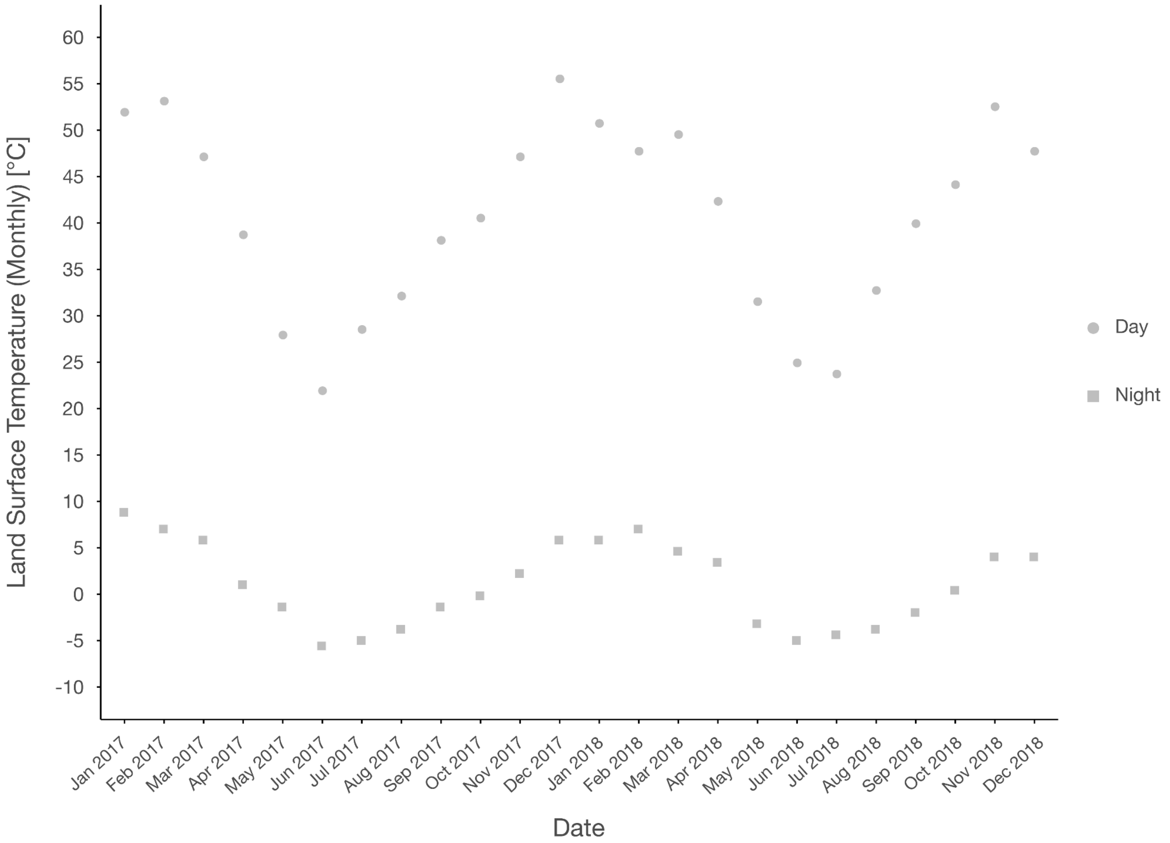


**Supplementary Figure S3. The monthly land surface temperature in the Laguna Pozo Bravo region shows great fluctuations between day and night as well as across seasons.** Monthly land surface temperature in degrees Celsius (°C) for the Laguna Pozo Bravo region from January 2017 to December 2018.

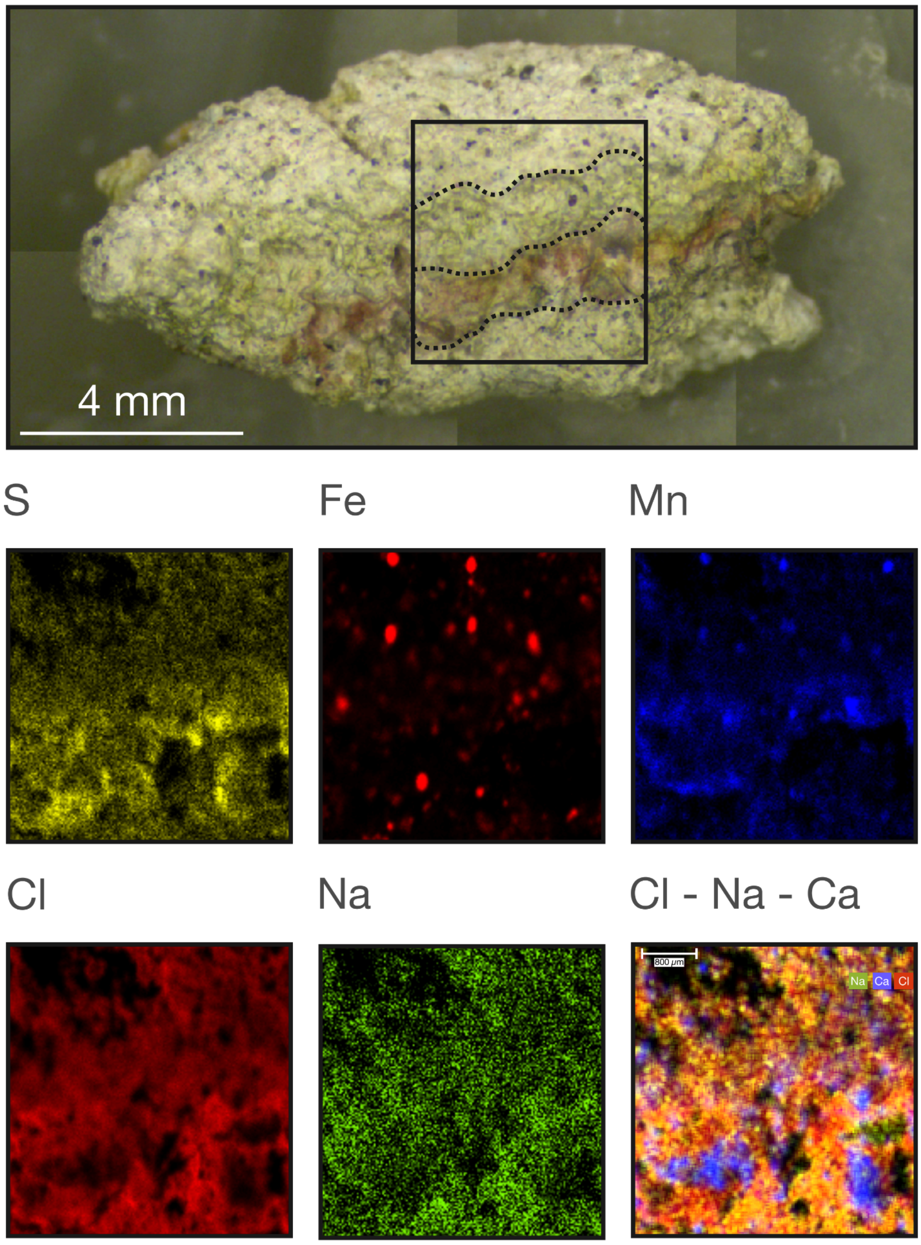


**Supplementary Figure S4. Pozo Bravo soft microbial mats show spatial heterogeneity in their chemical composition.** Spatial distribution of elements (deconvoluted counts) in a freeze-dried soft microbial mat sample determined by µXRF spectrometry. The area of µXRF scanning (4.2 mm x 4.3 mm) is marked with a black frame, and the dotted lines highlight the green and red/purple layers.

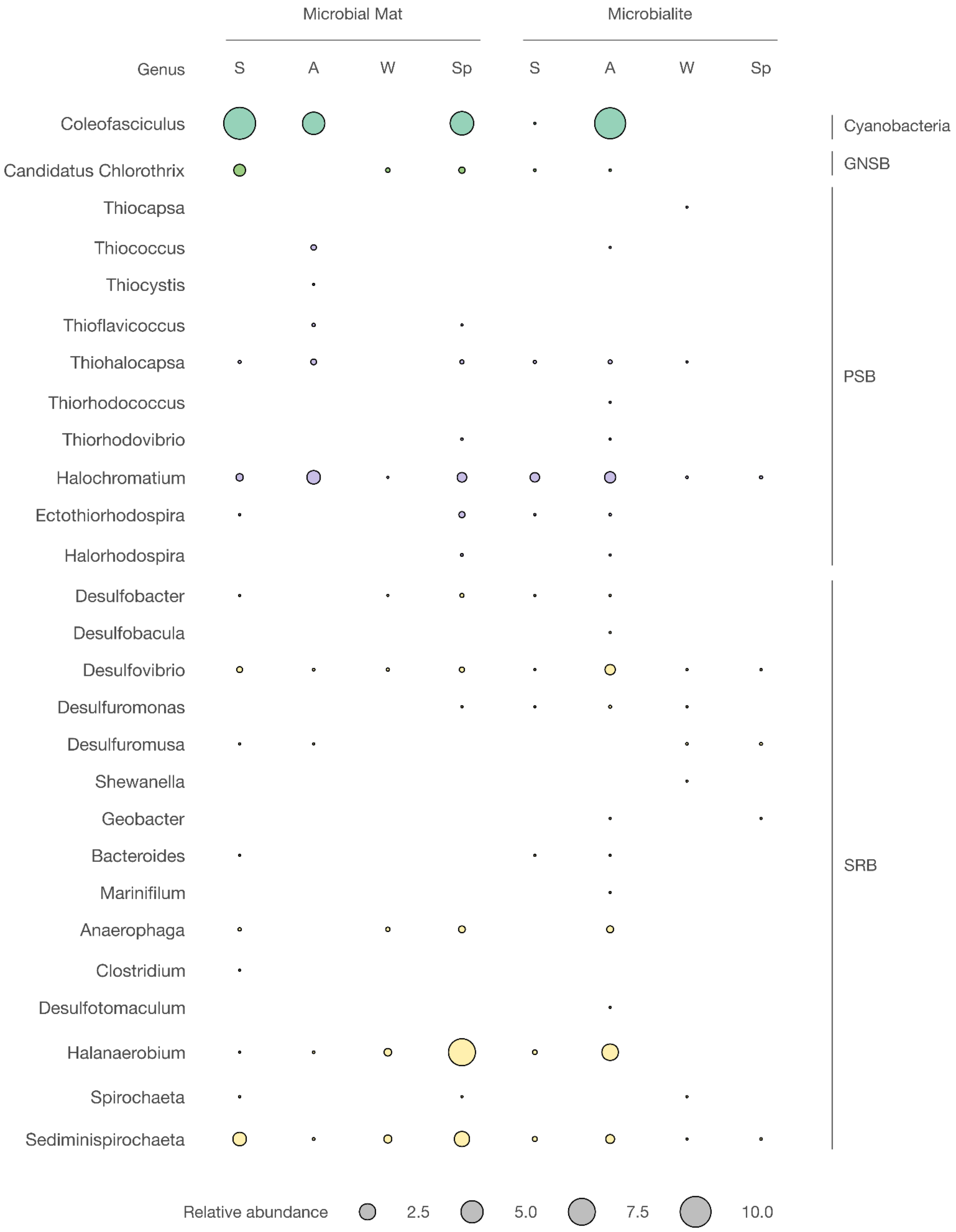


**Supplementary Figure S5. Seasonal shifts in the abundance of prokaryotic guilds within microbial mats and microbialites.** S, Summer; A, Autumn; W, Winter; Sp, Spring;

GNSB, Green non-sulphur bacteria; PSB, Purple sulphur bacteria; SRB, Sulphur reducing bacteria.

**Supplementary Tables**

**Supplementary Table S1**. Evaluation of elemental composition in the mapped area of a soft microbial mat sample. Standardless quantification for bulk material was performed with M4 Esprit software.

| Element | AN | Net counts | norm. C (wt.%) |
|---|---|---|---|
| Na | 11 | 75,122 | 27.23 |
| Mg | 12 | 18,583 | 1.79 |
| Al | 13 | 14,791 | 0.45 |
| Si | 14 | 829,153 | 9.9 |
| S | 16 | 264,444 | 1.13 |
| Cl | 17 | 7,998,234 | 37.08 |
| K | 19 | 107,907 | 0.67 |
| Ca | 20 | 5,133,361 | 20.06 |
| Ti | 22 | 45,620 | 0.1 |
| Mn | 25 | 368,049 | 0.35 |
| Fe | 26 | 1,326,688 | 0.97 |
| As | 33 | 151,041 | 0.05 |
| Sr | 38 | 566,336 | 0.21 |
| Rh | 45 | 304,933 | 0 |

**Supplementary Table S2**. Simulated saturation indices (SI) for Laguna Pozo Bravo during winter, using PHREEQC v3.0 geochemical software.

| Phase | SI | log IAP | log K (282 K, 1 atm) | Formula |
|---|---|---|---|---|
| Anhydrite | -0.88 | -4.93 | -4.05 | $CaSO_4$ |
| Aragonite | 2.78 | -5.35 | -8.13 | $CaCO_3$ |
| Arcanite | -4.74 | -6.87 | -2.13 | $K_2SO_4$ |
| Artinite | -1.75 | 19.20 | 20.95 | $Mg_2CO_3(OH)_2{\cdot}3H_2O$ |
| Bischofite | -6.24 | -1.45 | 4.79 | $MgCl_2{\cdot}6H_2O$ |
| Bloedite | -5.53 | -7.88 | -2.35 | $Na_2Mg(SO_4)_2{\cdot}4H_2O$ |
| Borax | -8.65 | 3.82 | 12.46 | $Na_2(B_4O_5(OH)_4){\cdot}8H_2O$ |
| Boric acid, s | -3.01 | -3.04 | -0.03 | $B(OH)_3$ |
| Brucite | -3.96 | -15.04 | -11.08 | $Mg(OH)_2$ |
| Burkeite | -8.19 | -8.97 | -0.77 | $Na_6CO_3(SO_4)_2$ |
| Calcite | 2.94 | -5.35 | -8.29 | $CaCO_3$ |
| Carnallite | -7.39 | -3.05 | 4.33 | $KMgCl_3{\cdot}6H_2O$ |
| $CO_2(g)$ | -1.20 | -2.45 | -1.25 | $CO_2$ |
| Dolomite | 6.07 | -10.64 | -16.71 | $CaMg(CO_3)_2$ |
| Epsomite | -3.17 | -5.15 | -1.98 | $MgSO_4{\cdot}7H_2O$ |
| Gaylussite | 0.60 | -8.82 | -9.42 | $CaNa_2(CO_3)_2{\cdot}5H_2O$ |
| Glaserite | -7.67 | -11.72 | -4.05 | $NaK_3(SO_4)_2$ |
| Glauberite | -2.53 | -7.78 | -5.25 | $Na_2Ca(SO_4)_2$ |
| Goergeyite | -1.06 | -31.56 | -30.50 | $K_2Ca_5(SO_4)_6{\cdot}H_2O$ |
| Gypsum | -0.41 | -5.01 | -4.61 | $CaSO_4{\cdot}2H_2O$ |
| $H_2O(g)$ | -1.98 | -0.04 | 1.94 | $H_2O$ |
| Halite | -1.13 | 0.41 | 1.54 | NaCl |
| Hexahydrite | -3.59 | -5.11 | -1.52 | $MgSO_4{\cdot}6H_2O$ |
| Huntite | 8.73 | 20.82 | 12.09 | $CaMg_3(CO_3)_4$ |
| $K_2B_4O_7{\cdot}4H_2O$ | -13.86 | 0.04 | 13.91 | $K_2B_4O_7{\cdot}4H_2O$ |

**Supplementary Table S3**. Simulated saturation indices (SI) for Laguna Pozo Bravo during summer, using PHREEQC v3.0 geochemical software.

| Phase | SI | log IAP | log K (282 K, 1 atm) | Formula |
|---|---|---|---|---|
| Anhydrite | -0.58 | -4.8 | -4.23 | $CaSO_4$ |
| Aragonite | 3.36 | -4.84 | -8.21 | $CaCO_3$ |
| Arcanite | -4.85 | -6.75 | -1.91 | $K_2SO_4$ |
| Artinite | 0.13 | 19.94 | 19.81 | $Mg_2CO_3(OH)_2 \cdot 3H_2O$ |
| Bischofite | -5.67 | -1.06 | 4.62 | $MgCl_2 \cdot 6H_2O$ |
| Bloedite | -5.24 | -7.58 | -2.35 | $Na_2Mg(SO_4)_2 \cdot 4H_2O$ |
| Borax | -3.61 | 8.85 | 12.46 | $Na_2(B_4O_5(OH)_4) \cdot 8H_2O$ |
| Boric acid, s | -1.91 | -1.94 | -0.03 | $B(OH)_3$ |
| Brucite | -2.63 | -13.54 | -10.9 | $Mg(OH)_2$ |
| Burkeite | -6.98 | -7.75 | -0.77 | $Na_6CO_3(SO_4)_2$ |
| Calcite | 3.54 | -4.84 | -8.39 | $CaCO_3$ |
| Carnallite | -6.81 | -2.4 | 4.42 | $KMgCl_3 \cdot 6H_2O$ |
| $CO_2$(g) | -1.32 | -2.76 | -1.44 | $CO_2$ |
| Dolomite | 7.36 | -9.67 | -17.04 | $CaMg(CO_3)_2$ |
| Epsomite | -3.32 | -5.19 | -1.86 | $MgSO_4 \cdot 7H_2O$ |
| Gaylussite | 1.67 | -7.75 | -9.42 | $CaNa_2(CO_3)_2 \cdot 5H_2O$ |
| Glaserite | -7.58 | -11.41 | -3.83 | $NaK_3(SO_4)_2$ |
| Glauberite | -2.04 | -7.37 | -5.33 | $Na_2Ca(SO_4)_2$ |
| Goergeyite | -1.34 | -30.81 | -29.48 | $K_2Ca_5(SO_4)_6 \cdot H_2O$ |
| Gypsum | -0.32 | -4.92 | -4.6 | $CaSO_4 \cdot 2H_2O$ |
| $H_2O$(g) | -1.61 | -0.06 | 1.55 | $H_2O$ |
| Halite | -0.83 | 0.75 | 1.58 | NaCl |
| Hexahydrite | -3.57 | -5.13 | -1.56 | $MgSO_4 \cdot 6H_2O$ |
| Huntite | 11.64 | 22.1 | 10.46 | $CaMg_3(CO_3)_4$ |
| $K_2B_4O_7 \cdot 4H_2O$ | -8.89 | 5.01 | 13.91 | $K_2B_4O_7 \cdot 4H_2O$ |

| | | | | |
|---|---|---|---|---|
| Kainite | -6.1 | -6.3 | -0.19 | $KMgClSO_4·3H_2O$ |
| Kalicinite | -3.22 | -13.16 | -9.94 | $KHCO_3$ |
| $KB_5O_8·4H_2O$ | -7.85 | -3.18 | 4.67 | $KB_5O_8·4H_2O$ |
| Kieserite | -4.61 | -4.84 | -0.23 | $MgSO_4·H_2O$ |
| Labile S | -4.38 | -10.06 | -5.67 | $Na_4Ca(SO_4)_3·2H_2O$ |
| Leonhardite | -4.13 | -5.01 | -0.89 | $MgSO_4·4H_2O$ |
| Leonite | -7.79 | -11.77 | -3.98 | $K_2Mg(SO_4)_2·4H_2O$ |
| Magnesite | 3.0 | -4.83 | -7.83 | $MgCO_3$ |
| MgCl2·2H2O | -15.55 | -0.83 | 14.72 | $MgCl_2·2H_2O$ |
| MgCl2·4H2O | -7.96 | -0.94 | 7.02 | $MgCl_2·4H_2O$ |
| Mirabilite | -1.82 | -3.15 | -1.33 | $Na_2SO_4·10H_2O$ |
| Misenite | -74.66 | -85.46 | -10.81 | $K_8H_6(SO_4)_7$ |
| NaB5O8·5H2O | -7.04 | -1.15 | 5.89 | $NaB_5O_8·5H_2O$ |
| NaBO2·4H2O | -3.23 | 6.34 | 9.57 | $NaBO_2·4H_2O$ |
| Nahcolite | -0.33 | -11.07 | -10.74 | $NaHCO_3$ |
| Natron | -2.36 | -3.19 | -0.82 | $Na_2CO_3·10H_2O$ |
| Nesquehonite | 0.17 | -5.0 | -5.17 | $MgCO_3·3H_2O$ |
| Pentahydrite | -3.79 | -5.07 | -1.28 | $MgSO_4·5H_2O$ |
| Pirssonite | 1.66 | -7.57 | -9.23 | $Na_2Ca(CO_3)_2·2H_2O$ |
| Polyhalite | -7.51 | -21.25 | -13.74 | $K_2MgCa_2(SO_4)_4·2H_2O$ |
| Portlandite | -8.36 | -13.55 | -5.19 | $Ca(OH)_2$ |
| Schoenite | -7.55 | -11.88 | -4.33 | $K_2Mg(SO_4)_2·6H_2O$ |
| Sylvite | -2.22 | -1.34 | 0.88 | $KCl$ |
| Syngenite | -5.22 | -11.61 | -6.39 | $K_2Ca(SO_4)_2·H_2O$ |
| Teepleite | -3.63 | 7.21 | 10.84 | $Na_2B(OH)_4Cl$ |
| Thenardite | -2.28 | -2.57 | -0.29 | $Na_2SO_4$ |
| Trona | -2.42 | -13.8 | -11.38 | $Na_3H(CO_3)_2·2H_2O$ |

**Supplementary Table S4**. Mineral composition of the microbial mat and microbialite samples, determined through powder X-ray diffraction. SM, Soft microbial mat; LM1; Lithified mat (external black rim); LM2, Lithified mat (light internal area); MI1, Microbialite (surface); MI2, Microbialite (external black rim); MI3, Microbialite (light internal area); MI4, Microbialite (core).

| Sample | SM | LM1 | LM2 | MI1 | MI2 | MI3 | MI4 |
|---|---|---|---|---|---|---|---|
| Albite intermediate (%) | - | - | - | - | 5.9 | - | - |
| Albite, calcian, ordered (%) | - | - | - | - | - | - | 6.1 |
| Albite, disordered (%) | - | 10.3 | - | - | - | - | - |
| Andesine (%) | 18.7 | - | - | - | - | 11.5 | - |
| Anorthite sodian (%) | - | - | 20.0 | - | - | - | - |
| Calcite (%) | - | - | - | 73.6 | 91.8 | - | 85.5 |
| Calcite, syn (%) | - | - | - | - | - | 84.9 | - |
| Halite, syn (%) | 8.9 | - | - | - | - | - | - |
| Hydrotalcite, syn (%) | - | - | - | 2.5 | - | - | 3.0 |
| Illite (%) | - | - | - | 2.5 | - | 1.9 | - |
| Illite-2M#1 [NR] (%) | - | - | - | - | - | - | 5.4 |
| Illite, 1M (%) | - | - | - | - | 1.1 | - | - |
| Labradorite (%) | - | - | - | 15.1 | - | - | - |
| Magnesium calcite, syn (%) | 67.2 | 83.3 | 75.3 | - | - | - | - |
| Quartz (%) | - | 6.4 | - | - | 1.2 | - | - |
| Quartz low, syn (%) | - | - | 4.7 | 6.3 | - | - | - |
| Quartz, low (%) | 5.3 | - | - | - | - | 1.8 | - |

**Supplementary Table S5**. Organic geochemical bulk parameters of a soft microbial mat and a microbialite. TOC, content of total organic carbon; TN, content of total nitrogen; $\delta^{13}C$, stable-carbon isotopic composition of organic carbon; $\delta^{15}N$, stable-nitrogen isotopic composition of total nitrogen.

| | Soft microbial mat | Microbialite |
|---|---|---|
| TOC (% dry weight) | 2.1 | 0.35 |
| $\delta^{13}C$ (‰) | -18.2 | -23.4 |
| TN (% dry weight) | 0.42 | n.d. |
| $\delta^{15}N$ (‰) | 0.4 | n.d. |
| C/N ratio | 5 | - |

n.d., means not detected.

**Supplementary Table S6**. Statistics for the sequencing, assembly, and annotation of the Pozo Bravo microbial mat metagenome. CDSs, coding sequences; OTUs, operational taxonomic units; ORFs, open reading frames.

| Parameter | Value |
|---|---|
| High-throughput sequencing | |
| Platform | Illumina MiSeq |
| Type of reads | paired end |
| Accession number | ERR3082345 |
| Number of reads | 8,232,105 (x2) |
| GC content of reads (%) | 52 |
| Read length (bp) | 35–251 |
| Annotation of metagenomic reads | |
| Number of identified CDSs | 8,367,672 |
| Number of functionally annotated CDSs | 3,741,958 |
| Number of prokaryotic OTUs | 286 |
| Assembly of metagenomic reads | |
| Total length (Mb) (contigs ≥ 500 bp) | 233.2 |
| Number of contigs | 237,161 |
| Maximum contig length (kb) | 83.2 |
| N50 (bp) | 955 |
| Percentage of mapped reads | 67.78 |
| Contig annotation | |
| Number of ORFs | 1,072,274 |

**Supplementary Table S7**. Statistics of the metagenome-assembled genomes (MAGs) from the Pozo Bravo microbial mat.

| MAG | Completeness (%) | Contamination (%) |
|---|---|---|
| 1 | 80.79 | 1.34 |
| 2 | 93.90 | 1.16 |
| 3 | 90.20 | 1.20 |
| 4 | 89.07 | 1.11 |
| 5 | 94.89 | 0.57 |
| 6 | 57.53 | 0.00 |
| 7 | 98.72 | 0.85 |
| 8 | 98.51 | 0.58 |
| 9 | 89.73 | 0.89 |
| 10 | 88.86 | 1.00 |
| 11 | 96.11 | 0.51 |
| 12 | 74.80 | 3.3 |
| 13 | 96.77 | 1.1 |

**Supplementary Table S8**. Taxonomic assignment of the metagenome-assembled genomes (MAGs) from the Pozo Bravo microbial mat.

| MAG | Phylum | Class | Order | Family | Genus |
|---|---|---|---|---|---|
| 1 | Bacteroidota | Rhodothermia | Rhodothermales | Salinibacteraceae | JAAAPH01 |
| 2 | Bacteroidota | Rhodothermia | Rhodothermales | Salinibacteraceae | Salisaeta |
| 3 | Bacteroidota | Bacteroidia | Cytophagales | Bernardetiaceae | |
| 4 | Bacteroidota | Bacteroidia | CAILMK01 | JACPUW01 | |
| 5 | Planctomycetota | Phycisphaerae | Phycisphaerales | UBA1924 | UBA1924 |
| 6 | Planctomycetota | Phycisphaerae | Phycisphaerales | UBA1924 | |
| 7 | Cyanobacteriota | Cyanobacteriia | Cyanobacteriales | Geitlerinemaceae | PCC-9228 |
| 8 | Cyanobacteriota | Cyanobacteriia | Cyanobacteriales | Geitlerinemaceae | |
| 9 | Cyanobacteriota | Cyanobacteriia | Cyanobacteriales | Rubidibacteraceae | Halothece |
| 10 | Cyanobacteriota | Cyanobacteriia | Cyanobacteriales | Rubidibacteraceae | Halothece |
| 11 | Pseudomonadota | Gammaproteobacteria | Thiohalospirales | Thiohalospiraceae | |
| 12 | Desulfobacterota | Desulfobacteria | Desulfobacterales | Desulfobacteraceae | JAIPEK01 |
| 13 | Desulfobacterota | Desulfobacteria | Desulfobacterales | JAABSP01 | |

**Supplementary Table S9**. List of profile hidden Markov models (HMMs) and their predefined score thresholds (Aramaki, T. *et a*l. 2020) used to annotate genes involved in metabolic pathways and cycles of interest.

| Gene name | Description | HMM ID | Score threshold |
|---|---|---|---|
| Oxygenic photosynthesis | | | |
| *psaA* | Photosystem I P700 chlorophyll a apoprotein A1 [EC:1.97.1.12] | K02689 | 913.90 |
| *psaB* | Photosystem I P700 chlorophyll a apoprotein A2 [EC:1.97.1.12] | K02690 | 786.83 |
| *psbA* | Photosystem II P680 reaction center D1 protein [EC:1.10.3.9] | K02703 | 245.97 |
| *psbD* | Photosystem II P680 reaction center D2 protein [EC:1.10.3.9] | K02706 | 379.17 |
| Anoxygenic photosynthesis | | | |
| *pufA* | Light-harvesting complex 1 alpha chain | K08926 | 49.83 |
| *pufB* | Light-harvesting complex 1 beta chain | K08927 | 51.67 |
| *pufH* (*puhA*) | Photosynthetic reaction center H subunit | K13991 | 261.17 |
| *pufL* | Photosynthetic reaction center L subunit | K08928 | 357.27 |
| *pufM* | Photosynthetic reaction center M subunit | K08929 | 384.50 |
| *pscA* | Photosystem P840 reaction center large subunit | K08940 | 1011.93 |
| (Bacterio)chlorophylls synthesis | | | |
| *acsF* (*chlE*) | Magnesium-protoporphyrin IX monomethyl ester (oxidative) cyclase [EC:1.14.13.81] | K04035 | 335.03 |
| *bchE* | Anaerobic magnesium-protoporphyrin IX monomethyl ester cyclase [EC:1.21.98.3] | K04034 | 440.43 |
| Nitrogen fixation | | | |

| | | | |
|---|---|---|---|
| *nifH* | Nitrogenase iron protein NifH | K02588 | 372.17 |
| *nifD* | Nitrogenase molybdenum-iron protein alpha chain [EC:1.18.6.1] | K02586 | 542.67 |
| *nifK* | Nitrogenase molybdenum-iron protein beta chain [EC:1.18.6.1] | K02591 | 401.70 |
| *vnfH* | Vanadium nitrogenase iron protein | K22899 | 558.90 |
| *vnfD* | Vanadium-dependent nitrogenase alpha chain [EC:1.18.6.2] | K22896 | 886.20 |
| *vnfK* | Vanadium-dependent nitrogenase beta chain [EC:1.18.6.2] | K22897 | 822.70 |
| *vnfG* | Vanadium nitrogenase delta subunit [EC:1.18.6.2] | K22898 | 150.97 |
| Carbon fixation | | | |
| *cbbL* | Ribulose-bisphosphate carboxylase large chain [EC:4.1.1.39] | K01601 | 319.20 |
| *cbbS* | Ribulose-bisphosphate carboxylase small chain [EC:4.1.1.39] | K01602 | 40.90 |
| *prkB* | Phosphoribulokinase [EC:2.7.1.19] | K00855 | 163.8 |
| *aclA* | ATP-citrate lyase alpha-subunit [EC:2.3.3.8] | K15230 | 902.47 |
| *aclB* | ATP-citrate lyase beta-subunit [EC:2.3.3.8] | K15231 | 516.33 |
| *ccl* | Citryl-CoA lyase [EC:4.1.3.34] | K15234 | 255.47 |
| *ccsA* | Citryl-CoA synthetase large subunit [EC:6.2.1.18] | K15232 | 643.9 |
| *ccsB* | Citryl-CoA synthetase small subunit | K15233 | 691.5 |
| *acsC* (*cdhE*) | Acetyl-CoA decarbonylase/synthase, CODH/ACS complex subunit gamma [EC:2.1.1.245] | K00197 | 393.83 |
| *acsD* (*cdhD*) | Acetyl-CoA decarbonylase/synthase, CODH/ACS complex subunit delta [EC:2.1.1.245] | K00194 | 220.97 |
| *mcl* | Malyl-CoA/(S)-citramalyl-CoA lyase [EC:4.1.3.24 4.1.3.25] | K08691 | 401.33 |

| | | | |
|---|---|---|---|
| *mcr* (bacteria) | Malonyl-CoA reductase / 3-hydroxypropionate dehydrogenase (NADP+) [EC:1.2.1.75 1.1.1.298] | K14468 | 1584.00 |
| *4-hblA* | 4-hydroxybutyrate---CoA ligase (AMP-forming) [EC:6.2.1.40] | K14466 | 863.07 |
| *mcr* (archaea) | Malonyl-CoA/succinyl-CoA reductase (NADPH) [EC:1.2.1.75 1.2.1.76] | K15017 | 695.97 |
| *4-hblC* | 4-hydroxybutyrate---CoA ligase (AMP-forming) [EC:6.2.1.40] | K14467 | 982.73 |
| Hydrogen metabolism | | | |
| *hmd* | 5,10-methenyltetrahydromethanopterin hydrogenase [EC:1.12.98.2] | K13942 | 575.00 |
| *hyaB* (*hybC*) | Hydrogenase large subunit [EC:1.12.99.6] | K06281 | 657.00 |
| *hyaA* (*hybO*) | Hydrogenase small subunit [EC:1.12.99.6] | K06282 | 329.93 |
| Sulphur metabolism | | | |
| *sqr* | Sulfide:quinone oxidoreductase [EC:1.8.5.4] | K17218 | 237.37 |
| *fccA* | Cytochrome subunit of sulfide dehydrogenase | K17230 | 79.87 |
| *fccB* | Sulfide dehydrogenase [flavocytochrome c] flavoprotein chain [EC:1.8.2.3] | K17229 | 372.00 |
| *soxA* | L-cysteine S-thiosulfotransferase [EC:2.8.5.2] | K17222 | 180.47 |
| *soxB* | S-sulfosulfanyl-L-cysteine sulfohydrolase [EC:3.1.6.20] | K17224 | 386.87 |
| *dsrA* | Dissimilatory sulfite reductase alpha subunit [EC:1.8.1.22] | K11180 | 307.20 |
| *dsrB* | Dissimilatory sulfite reductase beta subunit [EC:1.8.1.22] | K11181 | 355.17 |
| *hdrA* | Heterodisulfide reductase subunit A2 [EC:1.8.7.3 1.8.98.4 1.8.98.5 1.8.98.6] | K03388 | 517.40 |
| *hdrB* | Heterodisulfide reductase subunit B2 [EC:1.8.7.3 1.8.98.4 1.8.98.5 1.8.98.6] | K03389 | 320.83 |
| *sorA* | Sulfite dehydrogenase (cytochrome) subunit A [EC:1.8.2.1] | K05301 | 512.70 |

| | | | |
|---|---|---|---|
| *sorB* | Sulfite dehydrogenase (cytochrome) subunit B [EC:1.8.2.1] | K00386 | 85.93 |
| *soeA* | Sulfite dehydrogenase (quinone) subunit SoeA [EC:1.8.5.6] | K21307 | 437.03 |
| *soeB* | Sulfite dehydrogenase (quinone) subunit SoeB | K21308 | 227.30 |
| *aprA* | Adenylylsulfate reductase, subunit A [EC:1.8.99.2] | K00394 | 261.43 |
| *aprB* | Adenylylsulfate reductase, subunit B [EC:1.8.99.2] | K00395 | 79.70 |
| *sat* | Sulfate adenylyltransferase [EC:2.7.7.4] | K00958 | 271.63 |
| Nitrogen metabolism | | | |
| *tcdH* | Thiocyanate desulfurase [EC:1.8.2.7] | K24299 | 666.13 |
| *amoA* | Methane/ammonia monooxygenase subunit A [EC:1.14.18.3 1.14.99.39] | K10944 | 155.80 |
| *amoB* | Methane/ammonia monooxygenase subunit B | K10945 | 159.33 |
| *amoC* | Methane/ammonia monooxygenase subunit C | K10946 | 128.03 |
| *hao* | Hydroxylamine dehydrogenase [EC:1.7.2.6] | K10535 | 294.27 |
| *hzo* | Hydrazine dehydrogenase [EC:1.7.2.8] | K20935 | 1334.77 |
| *nxrA* | Nitrate reductase / nitrite oxidoreductase, alpha subunit [EC:1.7.5.1 1.7.99.-] | K00370 | 825.13 |
| *nxrB* | Nitrate reductase / nitrite oxidoreductase, beta subunit [EC:1.7.5.1 1.7.99.-] | K00371 | 341.60 |
| *narG* | Nitrate reductase / nitrite oxidoreductase, alpha subunit [EC:1.7.5.1 1.7.99.-] | K00370 | 825.13 |
| *napA* | Nitrate reductase (cytochrome) [EC:1.9.6.1] | K02567 | 806.27 |
| *nirK* | Nitrite reductase (NO-forming) [EC:1.7.2.1] | K00368 | 186.67 |
| *nirS* | Nitrite reductase (NO-forming) / hydroxylamine reductase [EC:1.7.2.1 1.7.99.1] | K15864 | 433.33 |

| | | | |
|---|---|---|---|
| *cnorB* | Nitric oxide reductase subunit B [EC:1.7.2.5] | K04561 | 136.13 |
| *qnorB* | Nitric oxide reductase subunit B [EC:1.7.2.5] | K04561 | 136.13 |
| *nosZI* | Nitrous-oxide reductase [EC:1.7.2.4] | K00376 | 468.97 |
| *nosZII* | Nitrous-oxide reductase [EC:1.7.2.4] | K00376 | 468.97 |
| Iron metabolism | | | |
| *cyc2* | Iron:rusticyanin reductase [EC:1.16.9.1] | K20150 | 888.27 |
| *rus* | Rusticyanin | K18683 | 256.80 |
| *mtrA* | Decaheme c-type cytochrome, DmsE family | TIGR03508.1 | 192.20 |
| *mtrB* | Decaheme-associated outer membrane protein, MtrB/PioB family | TIGR03509.1 | 240.05 |
| *mtrC* | Decaheme c-type cytochrome, OmcA/MtrC family | TIGR03507.1 | 209.30 |
| Arsenic metabolism | | | |
| *aioA* | Arsenite oxidase large subunit [EC:1.20.2.1 1.20.9.1] | K08356 | 671.93 |
| *aioB* | Arsenite oxidase small subunit [EC:1.20.2.1 1.20.9.1] | K08355 | 112.07 |
| *arrA* | Arsenate respiratory reductase molybdopterin-containing subunit ArrA | NF041714.1 | 1600.00 |
| *arrB* | Arsenate respiratory reductase iron-sulfur subunit ArrB | NF041715.1 | 450.00 |
| Manganese metabolism | | | |
| *moxA* | Manganese oxidase [EC:1.16.3.3] | K22348 | 370.83 |
| *cotA* | Spore coat protein A, manganese oxidase [EC:1.16.3.3] | K06324 | 554.50 |
| *mnxG* | Manganese oxidase [EC:1.16.3.3] | K22349 | 1132.33 |

| | | | |
|---|---|---|---|
| *mcoA* | Manganese oxidase [EC:1.16.3.3] | K22350 | 840.70 |
| Selenium metabolism | | | |
| *serA* | Selenate/chlorate reductase subunit alpha [EC:1.97.1.9 1.97.1.1] | K17050 | 2072.87 |
| *serB* | Selenate/chlorate reductase subunit beta [EC:1.97.1.9 1.97.1.1] | K17051 | 300.00 |

**Supplementary Table S10**. List of eukaryotic taxa identified in the Pozo Bravo microbial mat through metagenomic analysis.

| Phylum | Class | Order | Family | Genus | Description |
|---|---|---|---|---|---|
| Discosea | — | Longamoebia | Balamuthiidae | — | Free-living heterotrophic amoebae |
| Heterolobosea | — | Schizopyrenida | — | — | Heterotrophic amoeboflagellates |
| Euglenozoa | Kinetoplastea | Neobodonida | Rhynchomonadidae | Rhynchomonas | Heterotrophic kinetoplastid flagellates |
| | Dinophyceae | Syndiniales | — | — | Parasitic dinoflagellates |
| Ciliophora | Spirotrichea | — | — | — | Heterotrophic ciliates |
| Bacillariophyta | Bacillariophyceae | Naviculales | Phaeodactylaceae | Phaeodactylum | Oxygenic phototrophic diatoms |
| Bacillariophyta | Bacillariophyceae | Thalassiophysales | Catenulaceae | Amphora | Oxygenic phototrophic diatoms |
| Bacillariophyta | Bacillariophyceae | Bacillariales | Bacillariaceae | Nitzschia | Oxygenic phototrophic diatoms |
| Chlorophyta | Ulvophyceae | Ulotrichales | — | — | Oxygenic phototrophic green algae |
| Chlorophyta | Ulvophyceae | Ulvales | — | — | Oxygenic phototrophic green algae |

| | | | | | |
|---|---|---|---|---|---|
| Rhodophyta | Florideophyceae | Gracilariales | — | — | Oxygenic phototrophic red algae |
| Rhodophyta | Florideophyceae | Rhodymeniales | — | — | Oxygenic phototrophic red algae |
